# Accurate Inference of the Polyploid Continuum using Forward-time Simulations

**DOI:** 10.1101/2024.05.17.594724

**Authors:** Tamsen Dunn, Arun Sethuraman

## Abstract

Multiple rounds of whole-genome duplication (WGD) followed by diploidization have occurred throughout the evolutionary history of angiosperms. To understand how these cycles occur, much work has been done to model the genomic consequences and evolutionary significance of WGD. The machinations of diploidization are strongly influenced by the mode of speciation (allo or autopolyploidy). However, there is no discrete boundary between allo and autopolyploidy, which is best described as a continuum. Here we present a forward-time polyploid genome evolution simulator called SpecKS. SpecKS models polyploid speciation as originating from a 2D continuum, whose dimensions account for both the level of genetic differentiation between the ancestral parental genomes, as well the time lag between ancestral speciation and their subsequent reunion in the derived polyploid. Using extensive simulations, we demonstrate that changes in initial conditions along either dimension of the 2D continuum deterministically affect the shape of the *Ks* histogram. Our findings indicate that the error in the common method of estimating WGD time from the *Ks* histogram peak scales with the degree of allopolyploidy, and we present an alternative, accurate estimation method that is independent of the degree of allopolyploidy. Lastly, we use SpecKS to derive tests that infer both the lag time between parental divergence and WGD time, and the diversity of the ancestral species, from an input *Ks* histogram. We apply the latter test to transcriptomic data from over 200 species across the plant kingdom, the results of which are concordant with the prevailing theory that the majority of angiosperm lineages are derived from diverse parental genomes and may be of allopolyploid origin.

## Introduction

Multiple rounds of Whole-Genome Duplication (**WGD**) followed by diploidization have occurred throughout the evolutionary history of angiosperms (Otto and Whitton 2000; Soltis and Soltis 2012; Wendel 2015). WGD is considered a major speciation mechanism (Doyle and Egan 2010; Schranz et al. 2012; Wendel et al. 2016; Clark and Donoghue 2018), presenting a massive “macromutation”, potentially interfering with sexual reproduction, releasing transposons, and unbalancing molecular signaling pathways (McClintock 1929; Stebbins 1951; Mayer and Aguilera 1990; Comai et al. 2000; Ramsey and Schemske 2002; Le Comber et al. 2010; Arrigo and Barker 2012; Yant and Bomblies 2015; Bomblies et al. 2016; Zhang et al. 2016; Baduel et al. 2018). It has been theorized that the advantage of WGD lies not in the multiplicity of genomic material itself, but the intense period of genomic reorganization and gene shedding that follows, known as diploidization (Buggs et al. 2011; Madlung 2013; Soltis et al. 2014; Tank et al. 2015; Dodsworth et al. 2016; Soltis et al. 2016; Robertson et al. 2017; Baniaga et al. 2020; Carretero-Paulet and Van de Peer 2020; Nieto Feliner et al. 2020; Van de Peer et al. 2020; Li et al. 2021).

The diploidization process depends on whether an ancient WGD event is due to allopolyploidy (the multiplicity of genomes originating from two different species) or due to autopolyploidy (the multiplicity of genomes originating from within the same species). For example, allopolyploids are only capable of recombination across chromosomes from the same parental lineage, while autopolyploids are capable of recombination across all chromosomes (Blischak et al. 2023; Deb et al. 2023). Furthermore, the existence of intermediate types of polyploids suggests that polyploidy is best modeled as a continuum (Stebbins 1950; Meirmans and Van Tienderen 2013; Doyle and Sherman-Broyles 2017; Mason and Wendel 2020; Blischak et al. 2023).

The estimation of the relative proportion of speciation by allopolyploidy or autopolyploidy across the plant tree of life has been much debated (Stebbins 1947; Soltis et al. 2007; Soltis et al. 2010; Barker et al. 2016). Recent assessments suggest that the majority of sequenced plant genomes are derived from allopolyploidy (Darlington 1937; Clausen et al. 1945; Parisod et al. 2010; Soltis et al. 2010; Barker et al. 2016; Wang et al. 2019; Baniaga et al. 2020). Another critical parameter for assessing the evolutionary significance of WGD is the timing of the WGD event. This is critical for correlating WGD events with ecological and geological factors (Barba-Montoya et al. 2018; L. Zhang et al. 2020; Clark and Donoghue 2023), as well as understanding the timescale of the diploidization process (Hoegg et al. 2004; Doyle and Egan 2010; Mable 2013; De La Torre et al. 2017; Laurent et al. 2017; David et al. 2020; Li et al. 2021).

There are many methods to date WGD events. One method is to use genetic data to place the WGD event on a phylogenetic tree, if enough is known about its presence from multiple lineages (Bowers et al. 2003; Li et al. 2015; Li et al. 2018; Li and Barker 2020; Parey et al. 2022), or its presumptive parental species (Lott et al. 2009; Doyle and Egan 2010; Estep et al. 2014; Douglas et al. 2015; Thomas et al. 2017; Mccann et al. 2018; Wen et al. 2018; Yan et al. 2022; Conant 2023). If only the polyploid genome is available, a comparison of duplicated genes within the polyploid genome may be made to estimate their likely time of divergence (Lynch and Conery 2000; Blanc and Wolfe 2004; Cui et al. 2006; Vanneste et al. 2013; Clark et al. 2019; Chen and Zwaenepoel 2023). Syntenic relationships (self-synteny and with related species) may also be used to corroborate WGD (Vandepoele et al. 2002; Hampson et al. 2003; Wang et al. 2006; Parey et al. 2020).

Historically, the most common method to date a WGD event is to use a molecular clock to calibrate the divergence times between paralogs in a polyploid genome. The number of synonymous substitutions per synonymous site in protein-coding genes (***Ks***) is calculated between all paralogous pairs, and if there is a peak at *Ks>0,* the position of this peak is used to infer the time of WGD (Blanc and Wolfe 2004; Chen and Zwaenepoel 2023) and see figures 1 and 2.

**Figure 1.**
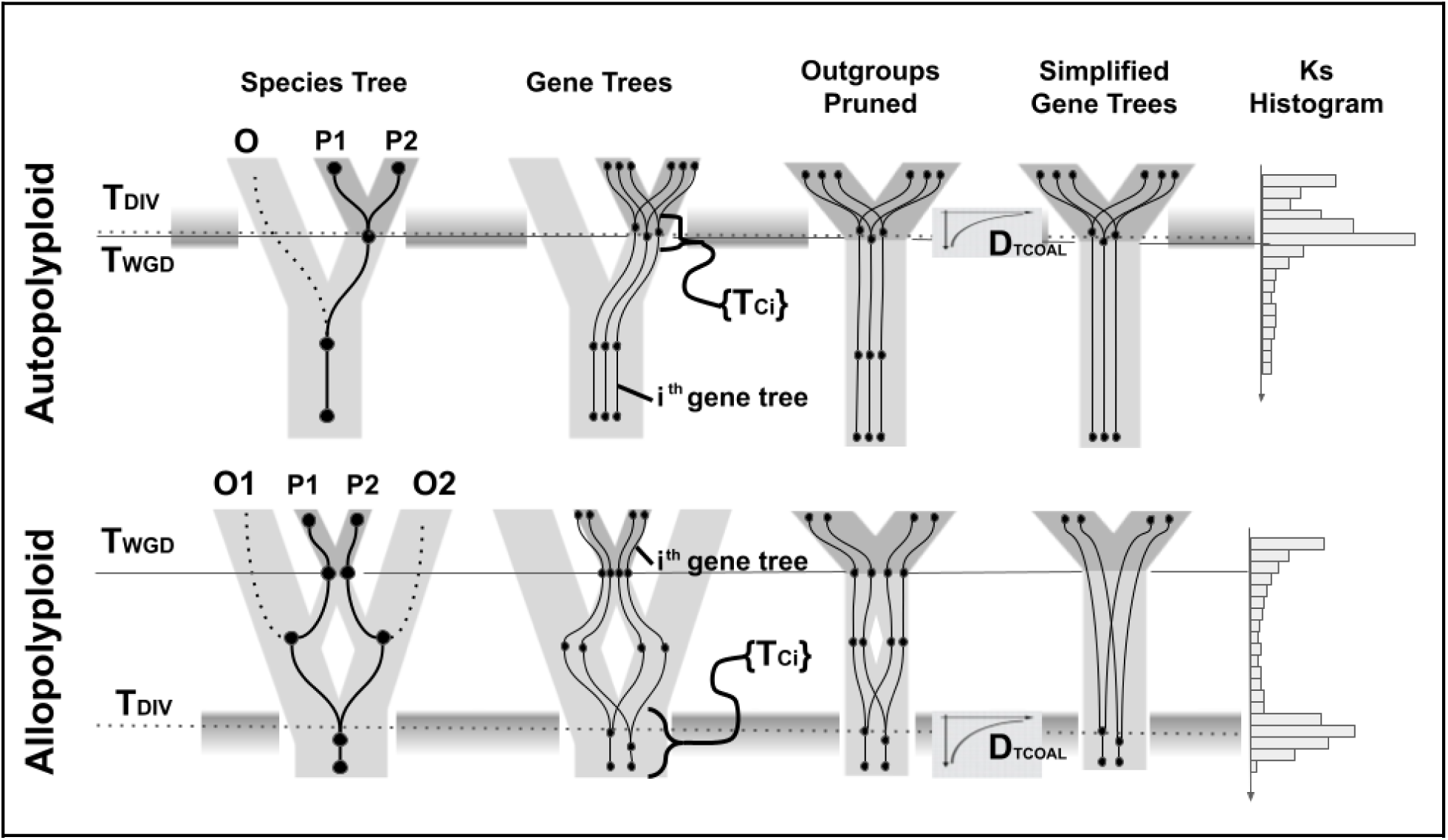
Relationship between the shape of the *Ks* histogram and the modes of auto versus allopolyploid speciation. Top: Autopolyploid speciation. Bottom: Allopolyploid speciation. Outgroup (O) and parental species are indicated in light gray. Polyploid species are shown in dark gray with duplicated genomes denoted as P1 and P2. The arrow in the *Ks* histogram points backwards in time. The set of gene tree coalescents {T_Ci_} are shown as nodes on the gene trees, and their distribution in time (D_TCOAL_) is indicated by the gray shaded bands. For allopolyploids the offset between T_WGD_ and T_DIV_ represents the lag time between separation of the diploid parental species (T_DIV_) and their later conjunction by polyploidization (T_WGD_). For autopolyploids, there may be instances where the gene tree divergence may begin before T_WGD_ or after it. In both cases, T_WGD_ and T_DIV_ are not synonymous, and D_TCOAL_ may be complex. Figures at the far right and left are derived from (Chen and Zwaenepoel, 2023).

**Figure 2.**
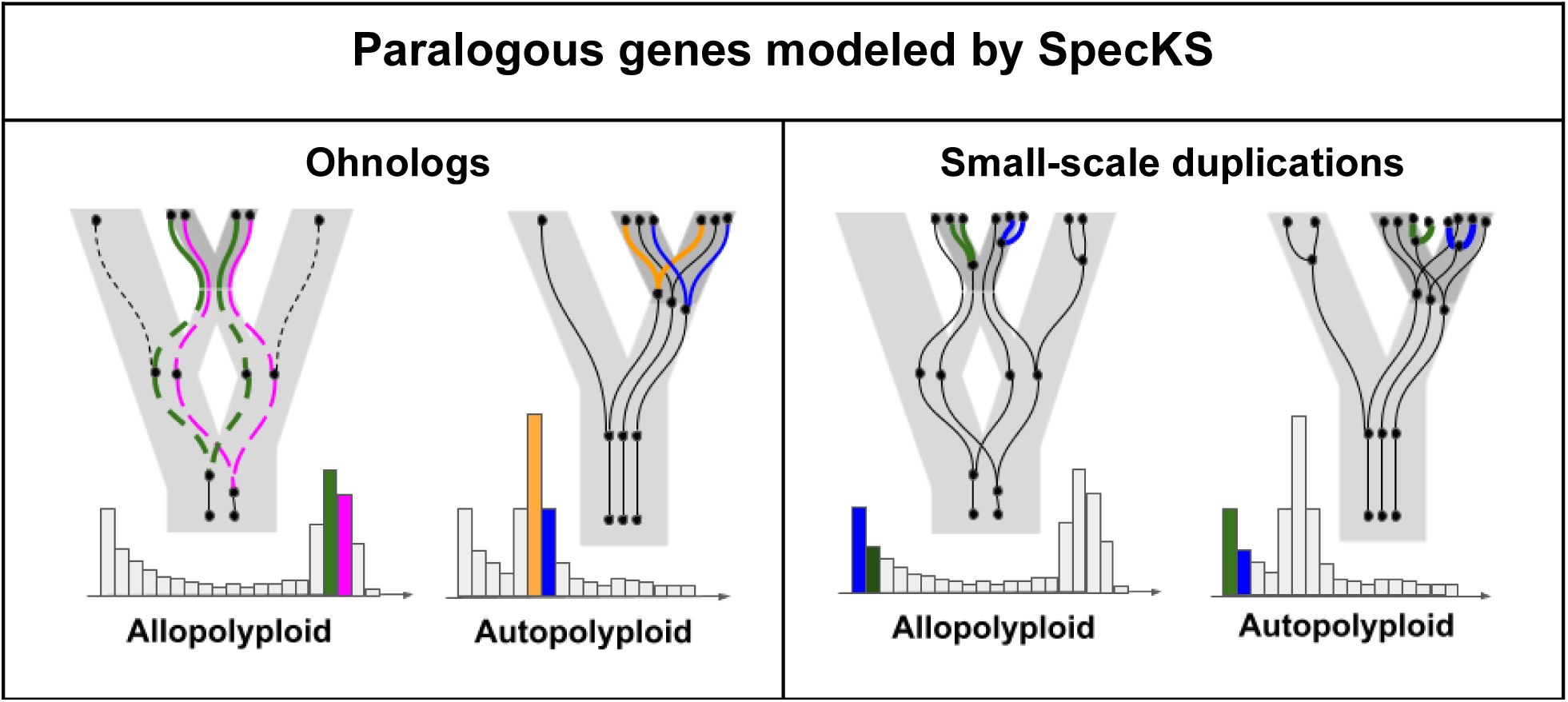
Ohnologs (paralogs generated by WGD) and small-scale duplications (“SSDs”, paralogs generated as single or small-scale copies) have unique contributions to the *Ks* histogram. Ohnologs (left panel) are shown for both allo and autopolyploids. Green and pink ohnologs (far left) are attributed to allopolyploidy and are born as orthologs (genes duplicated by speciation, dashed lines). Gold and blue ohnologs (center left) are attributed to autopolyploidy and are born by WGD. SSDs (right panel) follow the same birth and death process for both allo and autopolyploids and are shown here in green and blue. Unless maintained by selection, SSDs tend to be shed rapidly, and are thus found at the far left of the *Ks* histograms. As in Fig. 1, the polyploid species is shown in dark gray while the parental and outgroups are shown in light gray. The arrow of time points into the past along the X axis of the *Ks* histogram.

Since paralogs can arise from a number of evolutionary processes, *Ks* histogram shapes may be complex, and their interpretation may be challenging. *Ks* histograms generally have a first peak at *Ks=0*, due to the constant birth and death of small-scale duplications (SSDs). These SSD paralogs have recently arisen but have not yet been shed, and their lifespan follows an exponential decay model (Lynch and Conery 2000). If a species has undergone one or more WGD events, additional peaks will occur in the histogram for *Ks>0*, and these peaks may even be overlapping. These additional peaks are due to ohnologs (paralogs formed by WGD). The naive interpretation is that the mode in ohnolog divergence times represents the time of the WGD event (Ohno 1970; Blanc and Wolfe 2004). Univariate mixture models are used to empirically fit the peaks (Schlueter et al. 2004; Cui et al. 2006; Vanneste et al. 2013; Tiley et al. 2018; Li and Barker 2020; Chen and Zwaenepoel 2023).

However, the *Ks*-based approach has several pitfalls. The conversion between *Ks* and time is not straightforward (Wolfe et al. 1987; Doyle and Egan 2010; Barba-Montoya et al. 2018), *Ks* saturates for *Ks>2*, or roughly 200 million years (Cui et al. 2006; De La Torre et al. 2017; Li and Barker 2020), and multivariate fitting methods have been shown to overfit distributions (Vanneste et al. 2013; Tiley et al. 2018; Zwaenepoel and Van de Peer 2019). Restricting the *Ks* histogram to ohnologs may improve resolution (Van de Peer 2004; Zwaenepoel and Van de Peer 2019; Sensalari et al. 2022; Sutherland et al. 2024), but there remains significant methodological concerns.

While Ks-based methods can in theory correctly date the origin of a polyploid lineage which traces back to a single individual (assuming the genomes began to diverge at the same instant they duplicated (Doyle and Egan 2010)), the methods may fail for allopolyploids (Thomas et al. 2017; Mccann et al. 2018; Wen et al. 2018; Bouckaert et al. 2019; Conant 2023). This is because the peak of the *Ks* distribution corresponds to the divergence time between the diploid parental species (hereon **T_DIV_**), not the time of origin of the polyploid (hereon **T_WGD_**) (Doyle and Egan 2010; Thomas et al. 2017; Chen and Zwaenepoel 2023), as shown in figure 1. Since plant genomes can remain compatible for 10 mya or more after the last common ancestor (Senchina et al. 2003; Levin 2013) and up to 50 mya in one documented case (Rothfels et al. 2015), the difference between T_DIV_ and T_WGD_ can be significant. Confounding the *Ks* peak with T_WGD_ may also be problematic for autopolyploids, whose ohnologs may show complex patterns of divergence post WGD (Gaeta and Chris Pires 2010; Parey et al. 2022; Lv et al. 2024).

Several methods have been developed that can deal with the timing problems unique to allopolyploidy. These methods rely on additional data from the parental species or from broader sampling of the polyploid taxa (Thomas et al. 2017; Mccann et al. 2018; Wen et al. 2018; Bouckaert et al. 2019; Conant 2023). Population-demographic approaches have also been used with recent success to time WGD (Gutenkunst et al. 2009; St Onge et al. 2012; Roux and Pannell 2015; Roux et al. 2017; Blischak et al. 2023; Booker and Schrider 2023; Booker and Schrider 2023).

As part of the 1000 Plants (1KP) initiative (Leebens-Mack et al. 2019), transcriptomic data from over 1000 plants spanning the plant kingdom was used to compile *Ks* histograms. As in other works, Gaussian mixture models were fit and used to detect WGD from *Ks* peaks (Clark et al. 2019; Qiao et al. 2019; Guo et al. 2020; Li and Barker 2020). But a review of empirical *Ks* histogram shapes from the 1KP dataset show great complexity beyond the Gaussian (Zwaenepoel and Van de Peer 2019; Sensalari et al. 2022), which simulations have thus far have not been able to replicate (Sutherland et al. 2024).

Here we present the *Ks* simulator SpecKS. SpecKS simulates the forward evolution of polyploid genomes whereby the mode of speciation is not starkly allo or autopolyploid, but instead a point in a 2D continuum model where the parental species’ divergence in time and genetic space may vary independently (fig. 3), which may better align with our evolving definition of polyploidy (Parisod et al. 2010; Doyle and Sherman-Broyles 2017). Thus, SpecKS is highly configurable (tables S2 and S3) and capable of producing a rich array of *Ks* distribution shapes, modeling both the SSD and ohnolog components. Here we use SpecKS to demonstrate the sensitivity of the *Ks* distribution to several ancient speciation parameters, such as effective population sizes (N_e_), gene shedding rates, and the separation in time between T_DIV_ and T_WGD_.

**Figure 3.**
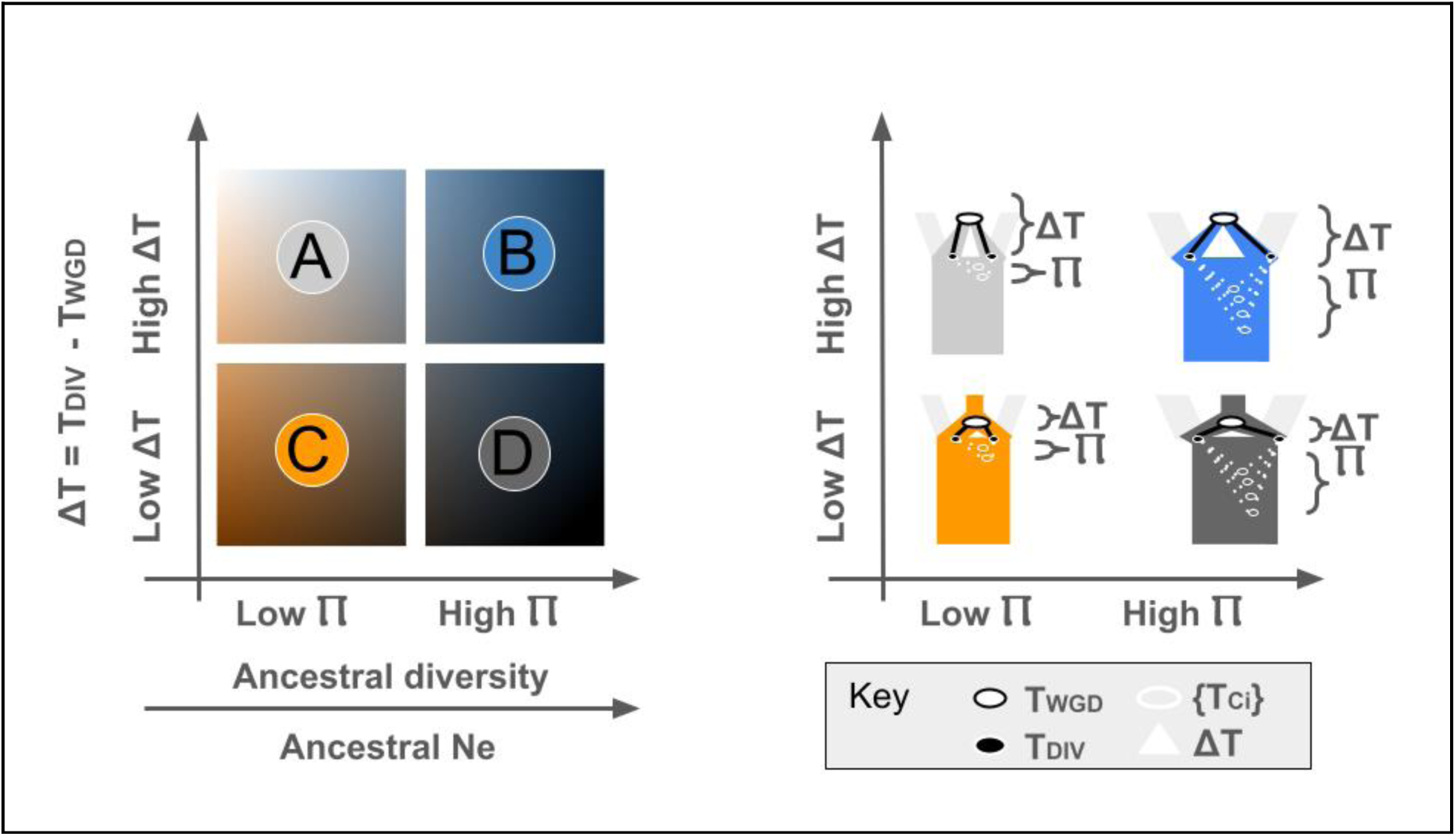
The SpecKS 2-dimensional model parameterizing allopolyploid speciation. A graphical representation of the 2D polyploid continuum modeled by SpecKS, showing Ⲡ, ΔT, T_DIV_, T_WGD,_ and the set of gene-tree coalescent times {T_COAL_i_} in the SpecKS model.

## Results

### Methods Overview

*Ks* histograms across the tree of life show a great diversity of distribution shapes, with some showing a high degree of symmetry while others appearing more skewed (Li and Barker 2020), suggesting the *Ks* histogram may bear signatures of evolutionary parameters or events beyond the presence or absence of WGD. Thus, we sought to build a simulation engine to investigate the potential effects of a variety of ancient polyploid speciation parameters on the distribution of *Ks* histogram shapes.

We developed a novel simulation-engine, SpecKS, which models polyploid speciation and evolution as a reticulate process. The simulation follows the evolution of an initial ancestral genome, which diverges at a given time (T_DIV_) into two sister diploid species. These sister species later recombine at T_WGD_, and the resulting polyploid continues to evolve to the present day. A set of *i* gene-trees {G_i_} are embedded in this reticulate topology with a configurable distribution (D_TCOAL_) of coalescent times {T_ci_}, which the user might base on ancestral diversity (Ⲡ), effective population size (N_e_), or some other relationship. Genes (as a set of random strings of nucleotides, {N_i_}) are evolved along {G_i_} using PAML v4.10.7 (Yang 2007) under neutrality to the present time. *Ks* is then calculated between the resulting paralogs, which are the leaves of the gene trees (fig. 4).

**Figure 4.**
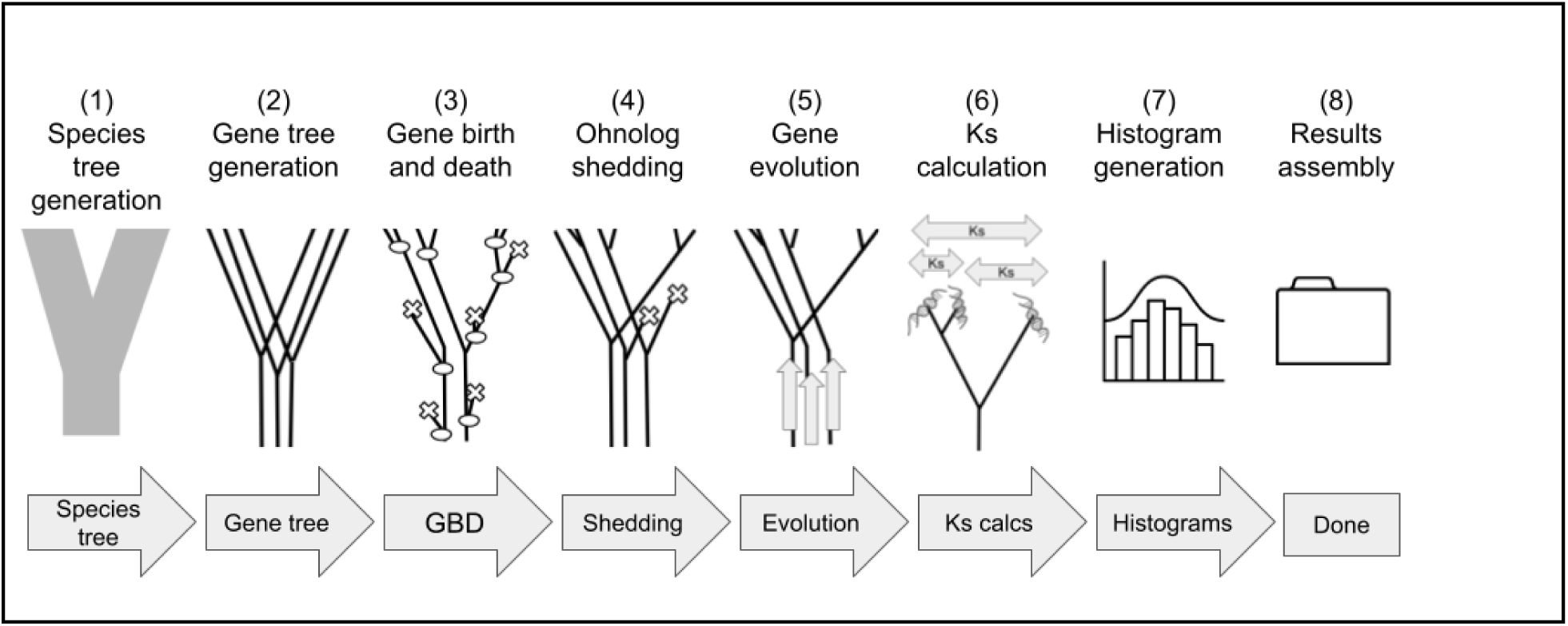
The SpecKS pipeline. A diagrammatic representation of the eight modules that make up the SpecKS polyploidy simulation pipeline.

Implicit in this model is the concept that the polyploid continuum has more than one dimension. We allow the degree of allopolyploidy to be a function of both length of the time the parental species were separated (ΔT) and the ancestral diversity (Ⲡ) of the subgenomes at T_DIV_. This allows us to separate the effects of these critical speciation parameters on the *Ks* histogram. We additionally disambiguate T_DIV_ and T_WGD_, which has confounded estimates of T_WGD_ in the past.

### SpecKS demonstrates that changes along either dimension of the 2D continuum will deterministically affect the shape of the *Ks* histogram

The initial distribution of the divergence times for the gene trees (D(T_c_)) is supplied by the user as an input parameter. This is to allow the user as much flexibility as possible with regard to modeling their system. To test if differences in these initial distribution shapes can be theoretically detected even after WGD and thus potentially affect the *Ks* histogram of extant polyploids, we simulated polyploids with modes of speciation from all four corners of our 2D continuum (fig. 3). Specifically, we tested sets of allopolyploids derived from (A) low-N_e_ ancestral species and a large ΔT between T_DIV_ and T_WGD_, (B) high-N_e_ ancestral species and a large ΔT, (C) low-N_e_ ancestral species and a small ΔT, and (D) high-N_e_ ancestral species and a small ΔT. Set (B) comprises canonical allopolyploids whose parental species were highly differentiated and have spent several million years apart before hybridization. Set (C) represents polyploids whose parental species were minimally differentiated and had no significant time between divergence and conjunction. The off-diagonals (A & D) represent plausible, but less intuitive polyploids. Specifically, set (A) are derived from parental species with little diversity between them at speciation, but a great amount of time between parental divergence and WGD. Set (D) are polyploids derived from parental species with a greater degree of diversity between them at speciation (highly differentiated subpopulations leading to separate species, but hybridization via polyploidy soon followed parental divergence). For all four sets, we simulated a range of WGD times, from ancient to relatively recent (80 - 0 MYA) (figure 5).

**Figure 5.**
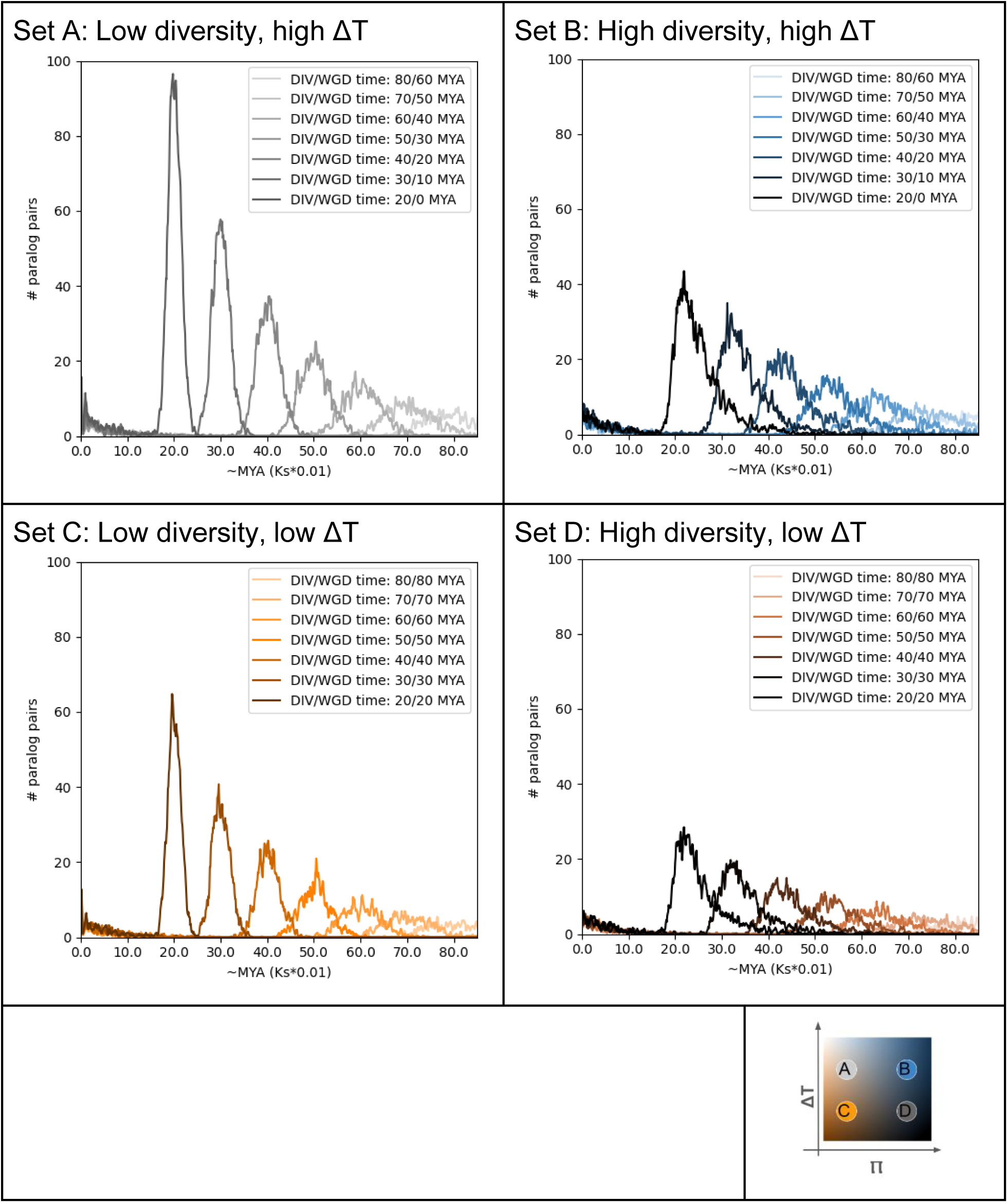
SpecKS demonstrates how the shapes of the *Ks* distributions attenuate over time. Set A: low Ne (1×10^5) ancestral species and a large ΔT (5 MY) Set B: high Ne (5×10^6) ancestral species and a large ΔT (5 MY) Set C: low Ne (1×10^5) ancestral species and a small ΔT (0 MY) Set D: high Ne (5×10^6) ancestral species and a small ΔT (0 MY)

With regard to the diversity dimension of the 2D continuum, our results showed that the polyploids derived from high-N_e_ ancestral species (A&B) show a more skew, fat-tailed WGD component in the *Ks* distribution compared to the low-N_e_ polyploids (figure 5). We also see that these differences persist for about 50 MY (figure 6, top). Along the ΔT dimension, we see that differences in ΔT had little effect on the skew but will affect the relative height differential between the SSD peak height and the WGD peak height (figure 5, and figure 6 bottom).

**Figure 6.**
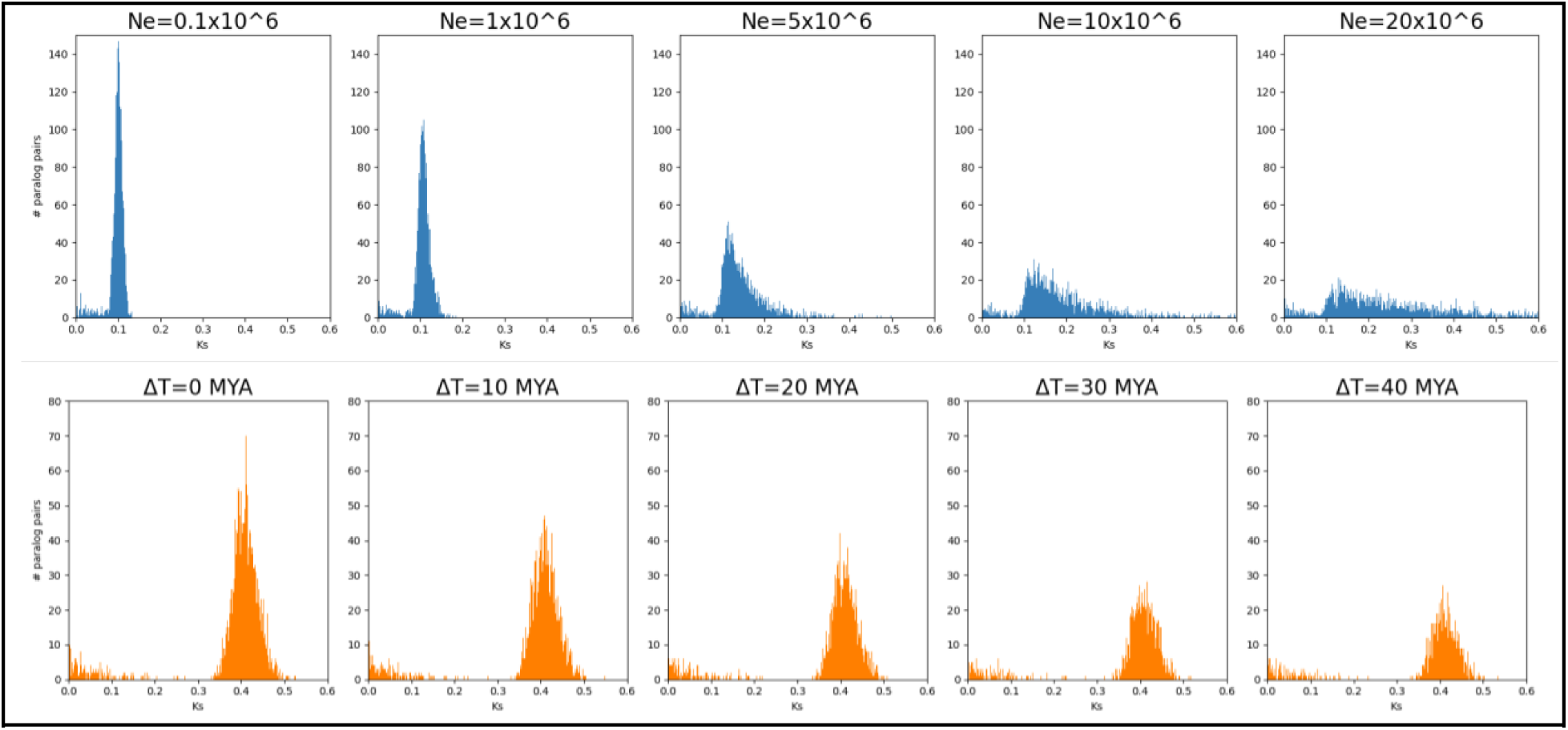
SpecKS demonstrates how the shapes of the *Ks* distribution vary with respect to ancestral diversity and ΔT. Top: Ancestral N_e_ varies from 10^5 to 20×10^6. T_WGD_ is fixed at 5 MYA and T_DIV_ is fixed at 10 MYA. Bottom: T_WGD_ varies from 0 to 40 MY, T_DIV_ is fixed at 40 MYA, and N_e_ is fixed at 1*10^6.

### The error in the common method of estimating WGD time from the *Ks* histogram peak scales with the degree of allopolyploidy

The “risk” of using the *Ks* peak to determine the timing of WGD, particularly for allopolyploids, has been well described by (Thomas et al. 2017; Chen and Zwaenepoel 2023). However, the *Ks* peak continues to be used to determine the timing of WGD. We were curious to see if SpecKS could be used to quantify the expected error in estimating T_WGD_, and potentially empirically relate the magnitude of T_WGD_ error to the degree of allopolyploidy along a continuum. We were further interested to test if an alternative method using the inferred start time of ohnolog-shedding would yield more accurate estimates of T_WGD_.

As expected, our simulations demonstrate that inferring T_WGD_ from the *Ks* peak may be quite problematic (off by millions of years). Since the *Ks* peak gives the T_DIV_, not the T_WGD_, it is no surprise that the error in estimates of T_WGD_ linearly scales with their difference. Thus, the error in inferring T_WGD_ from the *Ks* peak is proportional to ΔT, the degree of allopolyploidy as measured along the time axis of the 2D continuum (fig. 7, top).

**Figure 7.**
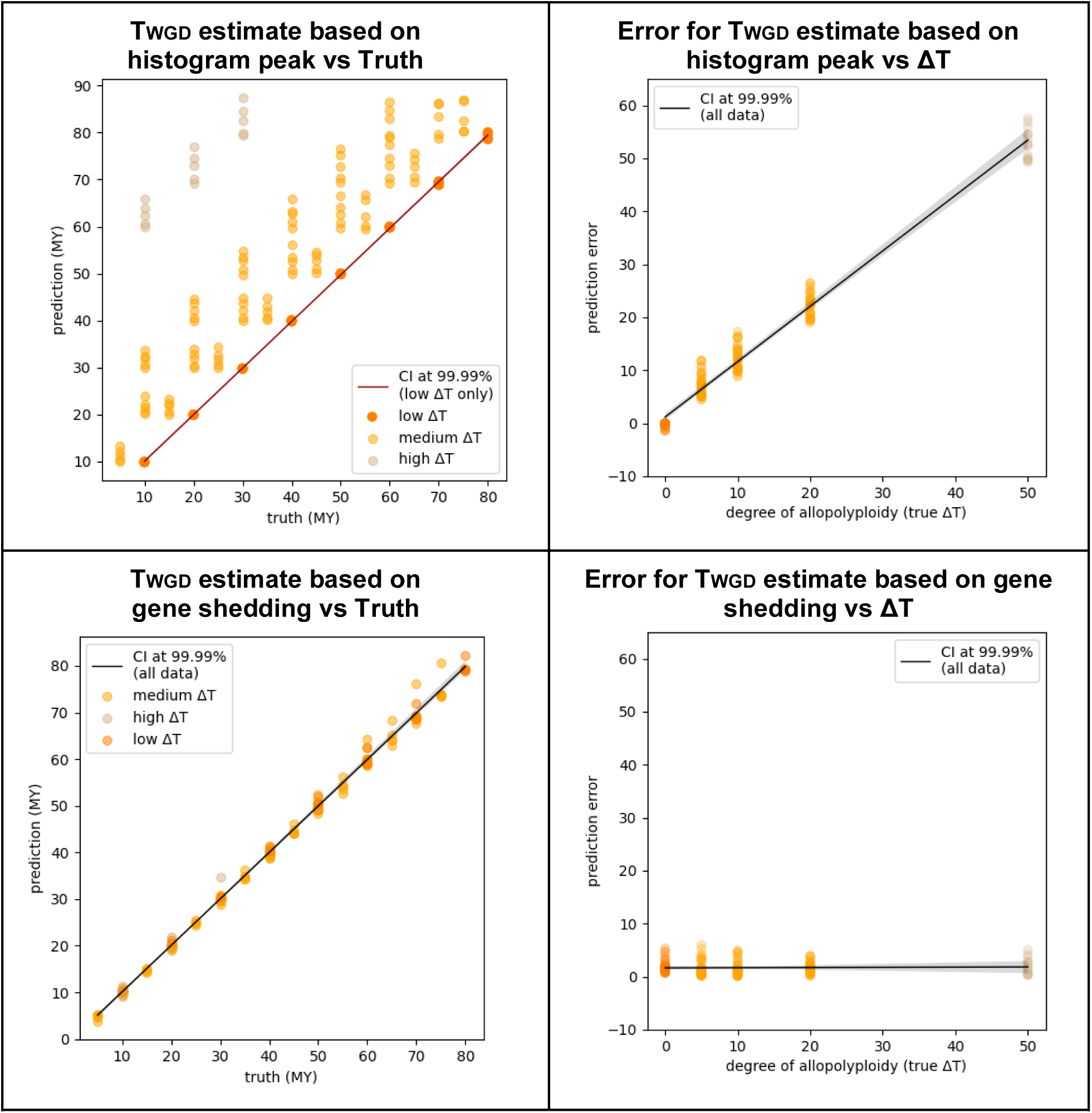
SpecKS shows that estimation of WGD time based number of genes remaining yields accurate results, irrespective of ΔT. (Left) T_WGD_ as estimated from the number of genes shed. (Top right) Error in the T_WGD_ as predicted from the number of genes shed (y axis) vs the degree of allopolyploidy (x-axis). (Bottom right) Error in T_WGD_ as estimated from the histogram mode (y axis) vs the degree of allopolyploidy (x-axis). Each datapoint represents results from simulations from dataset 1, described previously. Within dataset1, T_DIV_ times range from 10 to 80 MYA, by 10, with WGD offset (ΔT) by 0, 5, 10 and 50 MY. N_e_*G_t_ is 0.1,1.0,5.0, 10 and 20 MY.

### An alternative, accurate T_WGD_ estimation method that is independent of the degree of allopolyploidy

We tested the hypothesis that, since ohnolog-shedding can only begin after WGD irrespective of the mode of polyploid speciation, using the proportion of duplicate genes remaining may be a “less-risky” metric for inference, especially if the mode of speciation is unclear. In practice, the rates of gene shedding would vary by lineage and gene family, but for the purpose of this theoretical test, we assume the gene shedding rate is constant and known *a priori* for our theoretical species. We thus simulated a range of *Ks* histograms parameterized with a set gene shedding rate for a range of T_DIV_ and T_WGD_, for polyploids across a 2D polyploid continuum, thus recreating the range of possible *Ks* histograms for this hypothetical polyploid species, under a range of speciation scenarios. The simulations revealed a clear logarithmic relationship between the number of genes shed and T_WGD_, invariant to the mode of speciation (fig. S2, left). We were thus able to use the logarithmic relationship between T_WGD_ and the proportion of ohnologs remaining to determine T_WGD_ from the *Ks* histogram. In fig. 7, bottom left, we demonstrate the accuracy of this method, by comparing the input (true) T_WGD_ to the recovered T_WGD_, revealing a high accuracy, with r-value of 0.998 and a standard error of 0.004 MY. In fig. 7, bottom right, we show that the error (the difference between the true and inferred T_WGD_) does not increase with degree of allopolyploidy, as measured along the ΔT dimension.

### SpecKS recreates the main features of empirical *Ks* histograms

Next, we wanted to know if the evolutionary models incorporated in SpecKS would sufficiently reproduce the exponential decay curves typical of the primary “SSD” peak of the *Ks* histogram, as well as the secondary “WGD” peak, which contemporary works have had difficulty to replicate (Tiley et al. 2018; Sutherland et al. 2024). We selected three tetraploid species, *Coffea arabica, Zea mays* and *Populus trichocarpa* to demonstrate this. *Coffea arabica* is a relatively recent allopolyploid (WGD ∼10-600KYA). *Zea mays* was formed by polyploidy ∼14 MYA, and *Populus* ∼56 MYA (Gaut and Doebley 1997; Yu et al. 2011; Dai et al. 2014; Salojarvi et al. 2023). We show in figure 8 that SpecKS can well replicate both the SSD and ohnolog components of the *Ks* histograms for all three species. It is particularly interesting to note that the simulated *Ks* plots match the exponential-lognormal mixture models which have historically had the greatest success fitting observed *Ks* distributions. We note that this is an emergent property of our simulation, and no lognormal distributions were input. Furthermore, to achieve the best fit, the SpecKS input parameters were optimized to minimize the RMSE error (table S4), yielding T_WGD_ and T_DIV_ which correspond well with estimated dates from other sources (table S5) (Gaut and Doebley 1997; Yu et al. 2011; Dai et al. 2014; Salojärvi et al. 2024). We also note that the main difference between the simulated histograms and the true histograms is that the true histograms maintain a set of paralogs whose numbers do not decay over time and whose *Ks* distribution appears flat. One explanation for this phenomenon is that these paralogs are maintained by selection, and thus not presently modeled by our system.

**Figure 8.**
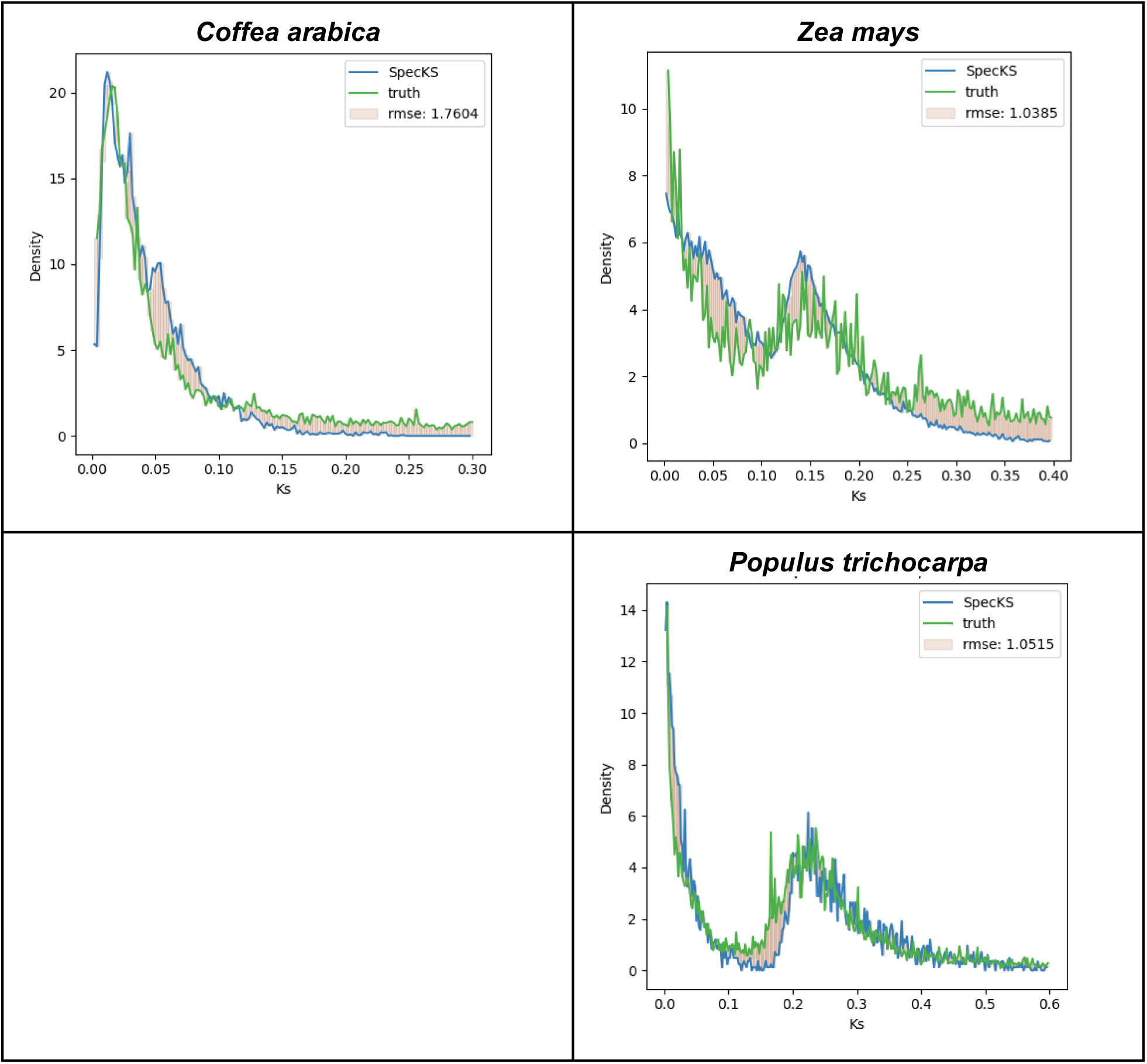
Using SpecKS to reproduce *Ks* empirical histograms from transcriptomic data. SpecKS-simulated histogram in blue and the empirical *Ks* histogram from transcriptomic data in green.

### Using SpecKS to estimate difference in time (ΔT=T_DIV_ -T_WGD_), and the diversity of the ancestral species (**Ⲡ**), from an input *Ks* histogram

As we demonstrated previously, the changes along either dimension of the 2D continuum deterministically affect the shape of the *Ks* histogram. Thus, we wanted to know if, given the *Ks* histogram, could the initial parameters of polyploid speciation be recovered? Since we have already demonstrated that T_WGD_ can be recovered (figure 7), what remains is to demonstrate the recovery of T_DIV_ and Ⲡ.

The peak of the *Ks* histogram has long been attributed to T_DIV_, albeit often confused with T_WGD_ (Blanc and Wolfe 2004), but disambiguated in (Thomas et al. 2017; Chen and Zwaenepoel 2023), thus we expected that the peak of the SpecKS-generated *Ks* histogram, once converted from *Ks*-space to time, SpecKS, should given an accurate recovery of the input T_DIV_.

Indeed, SpecKS corroborates this expectation (figure S2, right), and we demonstrate the accuracy of this method by comparing the input (true) T_DIV_ to the recovered T_DIV_, revealing a high accuracy, with r-value of 0.995 and a standard error of 0.008 MY. We note the caveat that this accuracy is contingent on the accuracy of the conversion factor between time and *Ks*, which is a configurable parameter in our simulation. Since ΔT = T_DIV_ -T_WGD_, the ability to estimate both T_DIV_ and T_WGD_ yields the estimated ΔT.

To test the recovery of ΔT, we generated a dataset of 160 simulations (dataset 1, parameters described in table 3 for a variety of modes of speciation across the 2D continuum, with a range of T_DIV_, T_WGD_, and Ⲡ. With these simulated datasets, using a ⅓ test, ⅔ training approach with the data, we trained a logistic regression classification model to discriminate between large, medium or small ΔT. Within the context of our simulated results, this approach gave 100% accuracy for WGD up to 80 MYA (figure 9).

**Figure 9.**
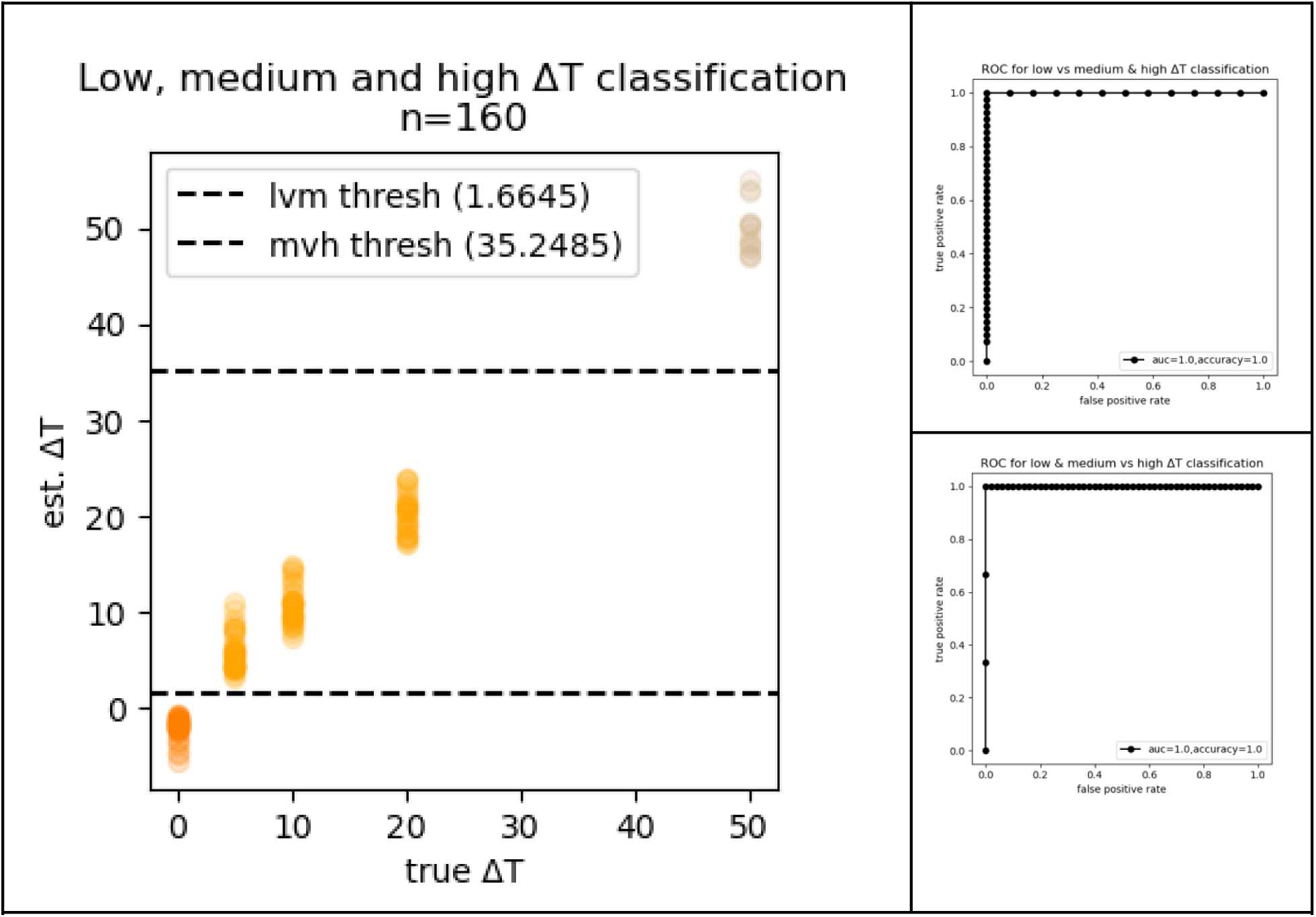
The low, medium and high ΔT discrimination model. (left) The ΔT discrimination threshold applied to the estimated ΔT. (right) Accuracy of inference of estimation ΔT, represented as an ROC plot. Each datapoint represents results from simulations from dataset 1, described previously. Lighter colors denote higher ΔT.

To test the recovery of Ⲡ, we chose a model (described in Methods) that would relate Ⲡ to the distribution of ortholog divergence times, since it is that distribution, which is input to SpecKS, not Ⲡ directly. Based on this model, we were able to use a range of N_e_ to generate a set of initial gene-tree divergence distributions as input to SpecKS. Using a ⅓ test, ⅔ training approach with the data, we trained a logistic regression classification model on dataset 1 to discriminate between high, medium or low ancestral N_e_. Within the context of our simulated results, this approach gave 94% accuracy for WGD up to 80 MYA (figure 10).

**Figure 10.**
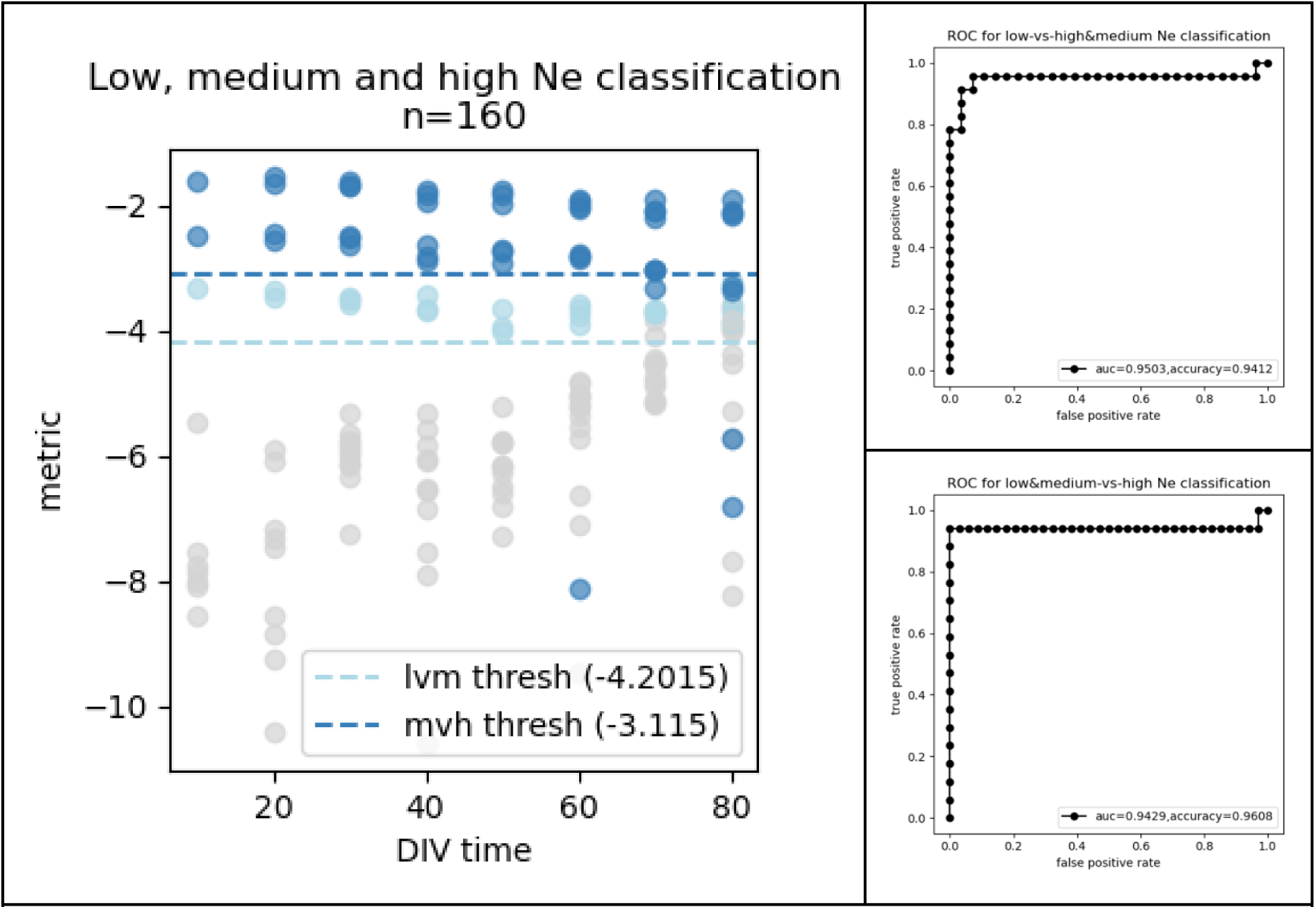
The low-vs-high Ne discrimination model. (left) The low-vs-high diversity discrimination threshold applied to simulated polyploids. The x-axis gives the SPC time. The y-axis gives the value of the high-vs-low diversity discrimination metric. (right) The ROC plot to assess accuracy of inference into high versus low Ne, across threshold values. Darker colors denote higher N_e_. Each datapoint represents results from a simulation from dataset 1, described previously. (Table 3).

### Using SpecKS to infer the polyploid continuum with *Ks* distributions from transcriptomic data from >200 angiosperms

Recent work has suggested that the majority of polyploid lineages may be derived from allopolyploidy rather than autopolyploidy (Wang et al. 2019), although also see (Soltis et al. 2007; Barker et al. 2016). Thus, we were interested to test if the logistical regression model discussed previously, when applied to empirical *Ks* data from species across the plant tree of life, would suggest high or low diversity between ancient subgenomes. Thus we applied our low-vs-high N_e_ classifier to over 200 real *Ks* observations made public by (Li and Barker 2020), derived from transcriptomic data from the 1KP study (Leebens-Mack et al. 2019). We were able to classify 228 WGD events, resulting in 12, 38 and 178, low-N_e_, medium-N_e_ and high-N_e_ determinations, respectively (fig. 11). Our results suggest that >94.7% of the WGD in the 1KP dataset are moderately or strongly allopolyploid, and the remaining 5.3% fall in an indeterminate category, with low diversity between the initial orthologs. These results are concordant with (Wang et al. 2019), suggesting that the majority of ancient polyploidization events which contributed to the genetic conduit may be derived from allopolyploidization with high ancestral N_e_. An alternative interpretation is that both our method and the method from (Wang et al. 2019) may under-report autopolyploids. One explanation for this might be that for autopolyploids, the gene copies have not diverged sufficiently to form true paralogs, thus the gene-pairs may not be detected as a multiplicity in transcriptomic analysis (Mayfield-Jones et al. 2013; Garsmeur et al. 2014). Furthermore, our logistical regression model was trained on simulations drawn from a continuum of allopolyploidy, thus may not be applicable to autopolyploids whose speciation parameters lie outside the scope of the SpecKS implementation. While much work exists describing the complexity of post-WGD divergence of ohnologs in specific autopolyploid systems, there currently exists no general, empirically-validated model (Parisod et al. 2010; Spoelhof et al. 2017; Parey et al. 2022; Lallemand et al. 2023). While the derivation of such a model is beyond the scope of this paper, the SpecKS architecture may prove to be well-suited to its incorporation and assessment.

**Figure 11.**
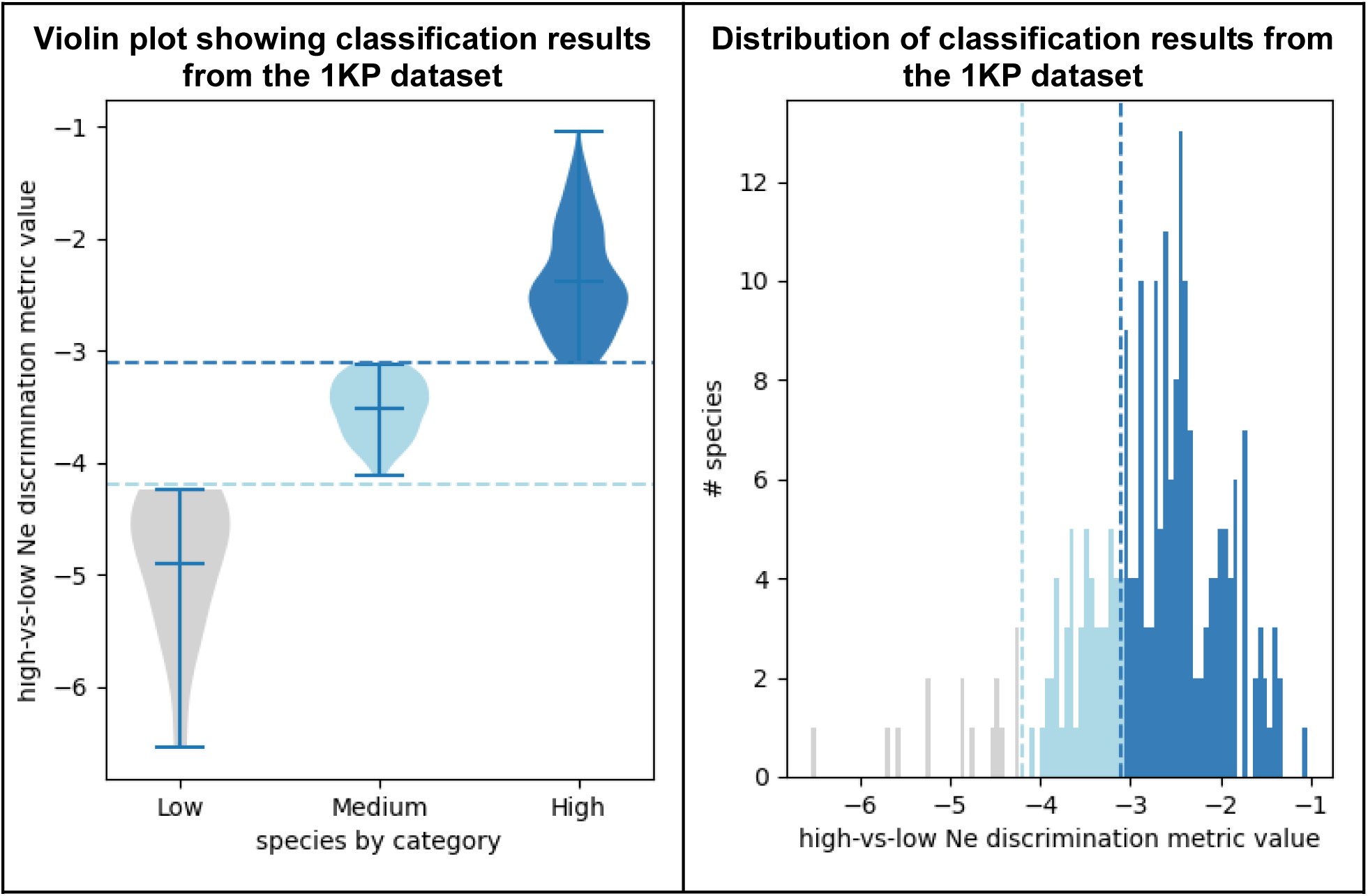
The Ne-classifier results for the 1KP dataset. (a) Violin plots indicating the number of species in each category. (b) Histogram giving the number of species in each category. Darker blue color represents higher-N_e_ ancestral species at SPC time, respectively. X-axis gives the numerical value of the discrimination metric. Y-axis gives the number of species per bin.

## Discussion

Here we present the polyploid genome evolution simulator SpecKS and use it to demonstrate the dependency of the shape of the Ks histogram on critical polyploid speciation parameters.

Our simulations also show that the shape and skew of the *Ks* histogram is sensitive to the evolutionary history of the ancestral polyploidization event: levels of ancestral genetic divergence, as well as the relative timing of T_WGD_ and T_DIV_, yielding characteristically different *Ks* distributions. We also demonstrate that these differences persist for 10s of millions of years (fig. 5). Furthermore, we have shown that the tail of the *Ks* histogram depends on the divergence between the ancestral subgenomes of the polyploid (fig. 6) and we demonstrate with simulations that the *Ks* distribution can be used to test inferences linking N_e_, Ⲡ, T_WGD_, T_DIV_, and distributions of ohnolog divergence times (fig. 9 and 10). Specifically, we demonstrate that the commonly used method for estimating T_WGD_ from the peak of the *Ks* histogram will be increasingly inaccurate for greater differences in time between T_WGD_ and T_DIV_ (fig. 7), and we develop an alternative method that works equally well for a variety of polyploid speciation scenarios. We contend that the *Ks* distribution is therefore information rich, with the potential to aid in the estimation of a variety of polyploid speciation parameters and scenarios.

We additionally show that SpecKS-generated histograms realistically capture the main features of empirical *Ks* histograms. For instance, in figure 8, we parameterized SpecKS to fit the observed *Ks* histograms of three well-studied species, coffee, maize and poplar. In all cases the histogram features corresponding to the SSDs and ohnolog peaks appear to be well replicated, with overall RMSE less than 2 in all cases (fig. 8). This demonstrates that the parameters supplied may be reasonably estimable under the SpecKS model. Interestingly, we note that the main difference between the simulated histograms and the true histograms, is that the true histograms maintain very old paralogous pairs (*Ks > T_DIV_*) whose numbers do not decay over time. In contrast, in our simulation, the number of paralogous pairs retained asymptotically approaches zero over time. The latter behavior is expected under the SpecKS model, because both SSDs and ohnologs are shed genes following a parameterized decay model (described in Methods).

One explanation for this phenomenon is that the paralogous pairs that persist in the true *Ks* histogram and not in the simulations may be duplicates that are maintained due to selection or potentially under linked selection (hitchhiking) and thus not presently modeled by our system. These persistent paralogs are more apparent in *Coffea arabica* and *Zea mays* (fig. 8). Since these species have undergone recent domestication (Gaut and Doebley 1997; Salojarvi et al. 2023; Yang et al. 2023), it may be possible that domestication has hindered efficient gene shedding or promoted paralog retention (Gaut et al. 2018). We also note that SpecKS currently only models ohnologs and SSDs; some paralogs of a different origin, like tandem duplications, may be retained for longer or shorter periods of time, and thus would be evident in the empirical histograms but not in the modeled histograms.

While SpecKS internal 2D continuum model is currently best suited to model varying degrees of allopolyploidy, it has not yet been extended to autopolyploidy. Furthermore, despite its utility, SpecKS currently represents a simplistic model which does not include selection, population dynamics, the effects of domestication, and other factors which would reasonably impact the *Ks* histogram. SpecKS’ internal model of evolution (GY94 with equal equilibrium codon frequencies) is simplistic, and the rate of *Ks* accumulation is assumed to be constant over time. Furthermore, our ability to test SpecKS against observed *Ks* histograms is limited to tetraploids whose WGD events are well-separated in time. This is because SpecKS does not yet model multiple superimposed rounds of WGD. An appropriate next step would be to iteratively include more evolutionary complexity, thus making the simulation both more realistic and more testable.

We note that many of the parameters incorporated into SpecKS are lineage specific (most significantly: rates of gene loss, the *Ks* distribution at the time of ancestral divergence, the rate of the molecular clock) and are difficult to ascertain *a priori*. Furthermore, the simplistic relationship between N_e_ and the gene tree coalescent distribution we selected as input (described in more detail in the Methods section) might not be appropriate for many use cases, including autopolyploids, or if complex population structure existed within the ancestral species. For these situations, we recommend the user consider the D_TCOAL_ most appropriate to their effort.

For simplicity, the discrimination models presented were drawn from simulation runs where a small number of parameters were fixed (for example, fixed gene-birth-and-death rate and molecular clock). These models are presented as proof of concept and not meant to analyze any particular lineage. When making lineage-specific inferences, we caution users to parameterize SpecKS with lineage-appropriate values, or if they are unknown, to use SpecKS over a range of likely values, to model the impact of uncertainty in their value.

## Materials and Methods

### Data

Transcriptomic data: *Ks* histograms in fig. 8 were generated from transcriptomic data for *Coffea arabic*a, *Zea mays* and *Populus trichocarpa*. Transcriptomic data were sourced from NCBI (NCBI assembly GCF_003713225.1, GCF_902167145.1 and GCF_000002775.5), and the software package KsRates (Sensalari, 2022) was used to calculate the *Ks* histograms for each species.

1KP data: In results section 8, the low-vs-high N_e_ discrimination model is run on histograms generated from *Ks* data from the 1KP dataset. The raw, empirical *Ks* data is available at (https://gitlab.com/barker-lab/1KP/-/tree/master/1KP_ksplots). These *Ks* data were generated by (Leebens-Mack et al. 2019; Li and Barker 2020). *Ks* histograms from this data were plotted using matplotlib (Hunter 2007), and the skew of the *Ks* histogram was measured using the same feature-extraction method used in the derivation of the N_e_ discrimination model, and then input directly to the N_e_ discrimination model.

### Logistic Regression Models

An example use-case for SpecKS is the provision of simulated truth and training data for the derivation of inference models. Here, we describe two discrimination models used in this paper, one to infer N_e_, and another to infer ΔT. For both of these models, we used the logistic regression package from scikit-learn (Pedregosa et al. 2011).

SpecKS-simulated *Ks* histograms were used to derive features, and SpecKS input parameters were used to derive target variables, which we organized into three categories (low, medium, and high) for each classifier. With respect to target variable categories, for the ΔT discrimination model, the target variable was based on whether ΔT=T_DIV_ -T_WGD_ < 5, < 30, or >= 30 MY. For the N_e_ discrimination model, the target variable was based on whether N_e_ < 5, < 10, or >= 10 * 10^6. These levels were selected to span realistic values (Gossmann et al. 2010). With respect to feature data, for the ΔT discrimination model, we used the difference between estimated T_WGD_ and T_DIV_ as our single feature, deriving both from the *Ks* histogram directly, using algorithms for estimating T_WGD_ and T_DIV_ as discussed previously in the results section (figures 7 and S2). For the N_e_ discrimination model, we used the natural logarithm of the skew of the *Ks* histogram (measured as the difference between the x-value of the *Ks* peak and the x-value of the *Ks* center of mass, in *Ks*-space) as the single key feature. The models were trained with a ⅓ test, ⅔ training approach, splitting datasets as appropriate.

### Setting the gene-tree divergence time distribution “D_TCOAL_”

In the SpecKS simulator, the initial distribution of the divergence times for the gene trees is supplied by the user as an input parameter. This allows the user as much flexibility as possible with regard to modeling their system. Throughout this paper, a simplistic input distribution D_TCOAL_ was used, which relates ancestral genetic diversity (Ⲡ) to the distribution of ortholog divergence times, based on the assumption of some level of allopolyploidy. We assumed that the ancestral diploid population was in panmixia, and we used the Kingman coalescent (Kingman 1982) to derive a diversity of initial gene-tree divergence times.

Under this model, the ancestral diploid species exists with some standing genetic variation which scales with population size, such that N_e_ (effective population size) is proportional Ⲡ. The ancestral species subsequently (at time T_DIV_) diverges to give rise to two diploid sister species, which evolve forward for a given amount of time ( ΔT = T_DIV_ -T_WGD_), before conjoining to form the diploid lineage at T_WGD_. Prior to T_DIV_, the coalescence times for gene copies between two random individuals in the ancestral population can be approximated by the Kingman coalescent, given Ⲡ. If these two ancestral individuals found new species, their individual sets of gene copies are now separated by speciation and are redefined as orthologs. The diversity of coalescent times becomes the initial diversity in nodes for the bifurcating gene trees at T_DIV_, which follows an exponential distribution, under the Kingman coalescent (Kingman 1982).

Under this simple one-genome-one-species model, for more diverse ancestral species, we see more initial diversity in node times, and a greater skew in the *Ks* distribution towards the past. Mathematically, this is because the decay constant in the Kingman coalescent is inversely proportional to N_e_. Note that this model may not be appropriate for autopolyploids, which is beyond the scope of this paper.

To obtain the Kingman coalescent from N_e_, it is necessary to assume a generation time (G_t_). In all our simulations, for simplicity, we assumed at G_t_=1. Since the *Ks* distributions generated and the inferences made from them were done in *Ks* space, the exact G_t_ is immaterial. It only matters that the Ks-to-My conversion factor, which is a configurable input, properly factors in the G_t_ at the initiation of the simulation, and that this same conversion factor is taken into account by the end-user when relating the output *Ks* plots to chronological time.

### SpecKS Implementation

SpecKS is implemented as a pipeline application in python. SpecKS takes as input an XML configuration file listing the simulation parameters (Table 1, S2 & S3) and outputs a text file (.csv) of all pairwise *Ks* accumulated between all gene pairs (ohnologs and small-scale duplications) for each simulated genome. Architecturally, SpecKS is designed as a pipeline with eight modules which are executed sequentially for each polyploid in the simulation (fig. 4). The modules functions are (1) species tree generation, (2) gene tree generation, (3) application of a gene birth and death model, (4) application of a post WGD gene shedding model, (5) gene sequence evolution, (6) the *Ks* calculation between gene pairs, (7) *Ks* histogrammer and (8) final results assembly. We give details for each module below.

1. Species tree generation: A newick-formatted species tree is generated for each polyploid. This is aways in the format (O:T_SIM_, (P1:T_DIV_, P2:T_DIV_): T_SIM_-T_DIV_), where T_SIM_ specifies the full simulation time. P1 and P2 denote the subgenomes of the polyploid, while O denotes the outgroup.
2. Gene tree generation: Newick-formatted gene trees are generated from the simulated species tree. Random variations in gene divergence times are introduced according to the distribution specified in the configuration file, allowing for the introduction of a range of divergence times for the ohnologs (genes duplicated by whole-genome duplication). The distribution is configurable and might in theory be selected based on the ancestral genetic diversity (Ⲡ), generation time, and evolutionary model. The number of gene trees is also configurable, with a default set at 3000. The distribution selected for our simulations was based on the Kingman coalescent, and we give details in the methods section.
3. Gene birth and death model: The gene birth and death model introduces small-scale duplications (“SSDs”) into the simulation. For each gene tree, genes are randomly born at a rate specified in the configuration file, modeled as a Poisson process (Zhao et al. 2015). Genes are assigned a death-date at birth, with a life span drawn randomly from an exponential decay distribution (Lynch and Conery 2000; Lynch et al. 2001). Gene birth rate and mean life expectancy are configurable (Lynch and Conery 2003; Guo 2013). Genes born which will be dead before the simulation concludes are pruned at this time for computational efficiency.
4. Ohnolog gene shedding model: Genes duplicated by WGD also have a mean life expectancy given by an exponential decay distribution (Ren et al. 2018). The proportion of WGD duplicates which will be dead before the simulation concludes are calculated based on WGD time and pruned for computational efficiency. For simplicity and traceability, ohnolog duplicates are always removed from the P1 branch. Ohnolog life expectancy is configurable and based on (Maere et al. 2005; Guo 2013).
5. Gene sequence evolution: For each simulated gene tree, we simulated codon sequences using the EVOLVER program within PAML v4.10.7 (Yang and Nielsen 2000: 200; Yang 2007). PAML was configured as in (Tiley et al. 2018) to simulate codon evolution along each finalized gene tree using a Goldman–Yang (GY94) model of codon evolution (Goldman and Yang 1994; Yang and Nielsen 1998) with equal equilibrium codon frequencies, a transition/transversion rate ratio of 2 and a global dN/dS of 0.2 as in (Tiley et al. 2018). The number of codons per alignment is set by default to 1000 as in (Tiley et al. 2018), in line with reported plant transcript lengths (Luo et al. 2019; S. Zhang et al. 2020; Al-Dossary et al. 2023), but is configurable.
6. *Ks* calculation: *Ks* is calculated between all pairs of terminal genes in a given gene tree, across all gene trees, using the CODEML program within PAML v4.10.7. Codon frequencies, transition/transversion rate and *dN/dS* were set to match those used in EVOLVER (Yang and Nielsen 2000; Yang 2007). *Ks* per million years is 0.01 by default and is configurable (Blanc and Wolfe 2004; Tiley et al. 2018).
7. *Ks* histogrammer: SpecKS thereon generates two default histograms (one with a configurable maximum *Ks*, and one with no maximum *Ks*). The existence of these histograms is only meant to give a confirmation of the run success. It is expected that the end users will use their own visualization tools to generate more elegant histograms.
8. Final results assembly: The final results assembler collates the CODEML results, producing a single .csv file containing the *Ks* results for each polyploid. Additional products of the simulation include *Kn* results (histogram of accumulated non-synonymous mutations between paralogs) for the polyploid, and *Ks* and *Kn* results for the outgroup.

**Table 1.**
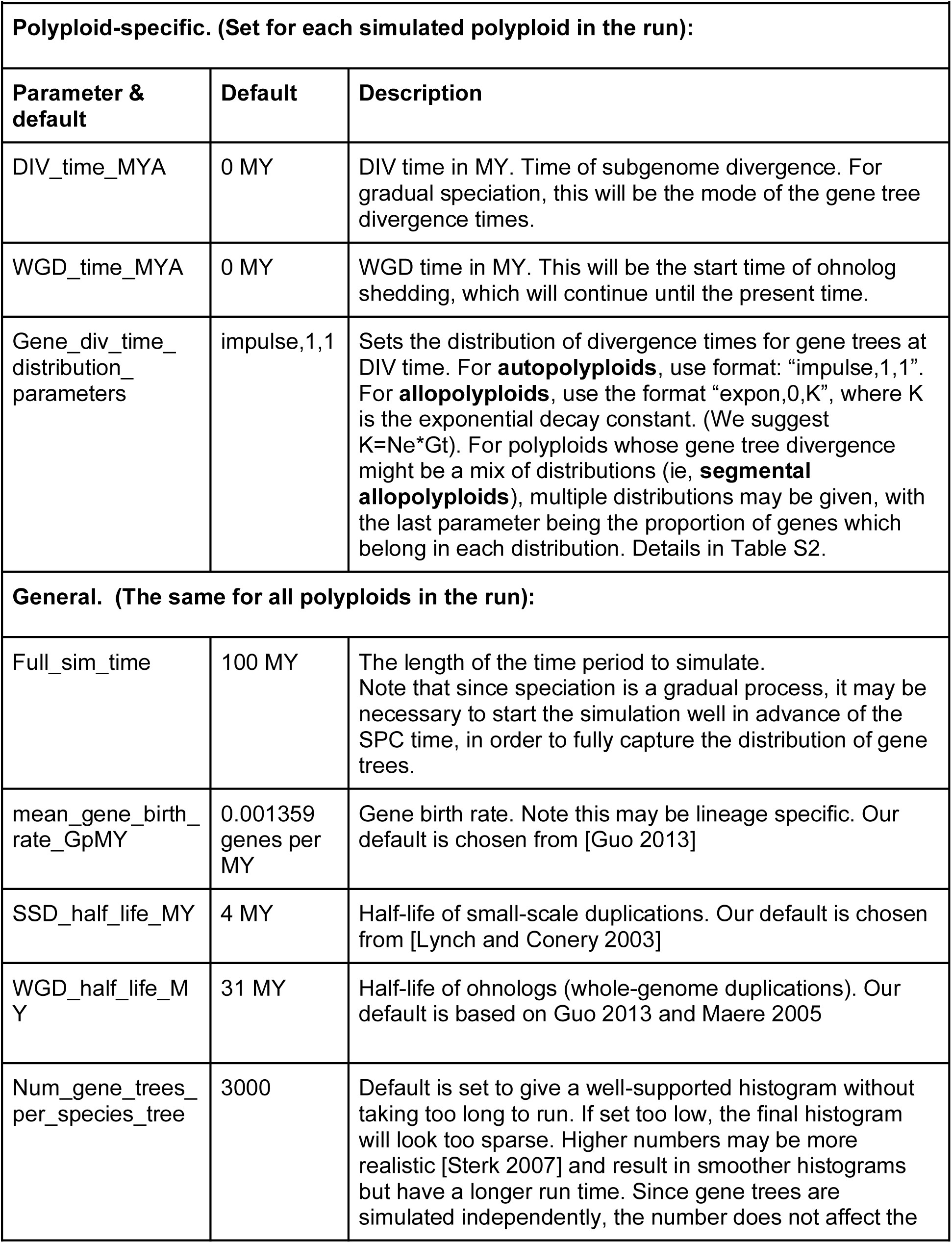
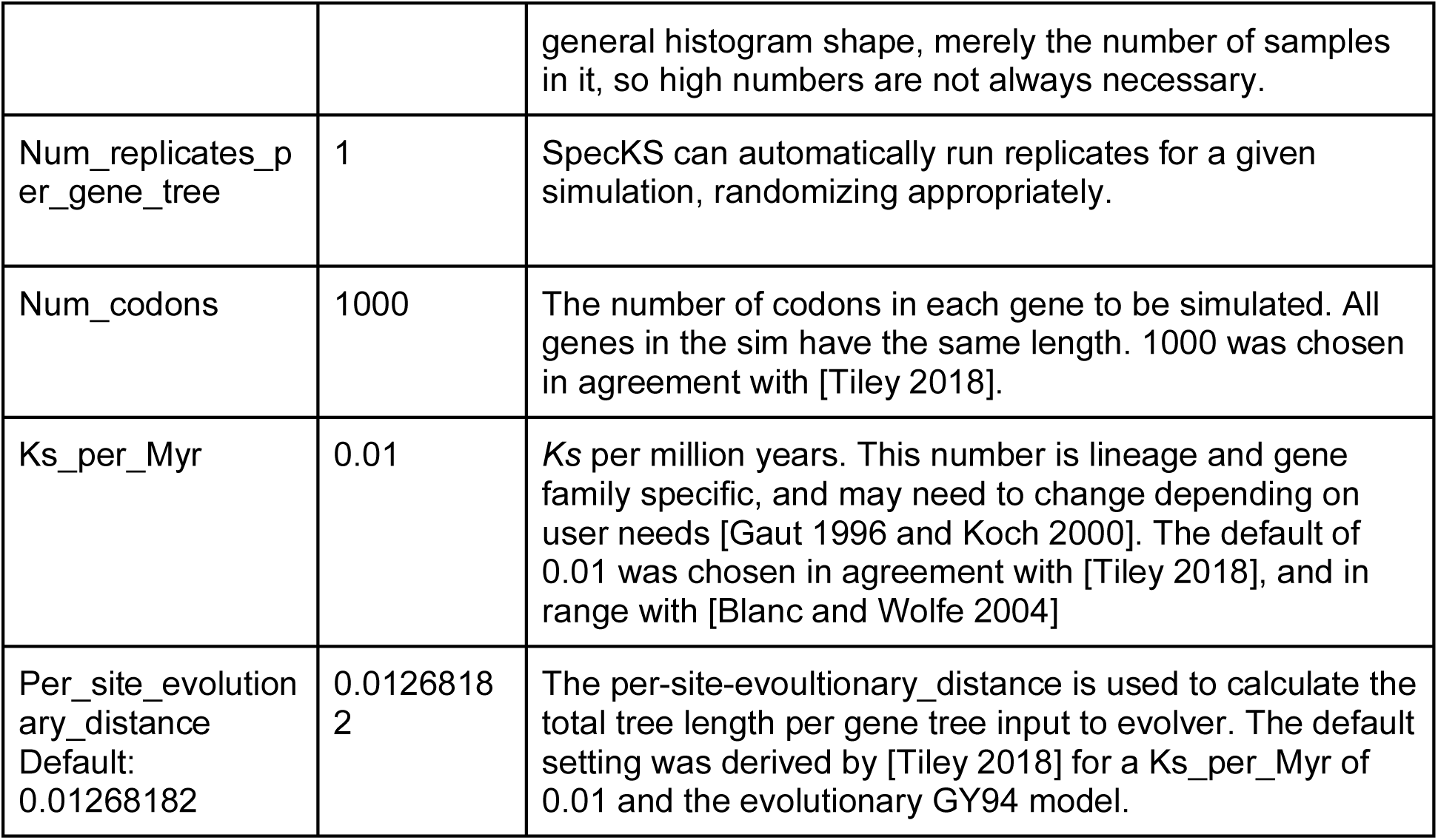
Subset of SpecKS configurable parameters.

**Table 2.** Summary of 1KP categorization results.

| **Table 2. Summary of 1KP categorization results** |  |  |  |
| --- | --- | --- | --- |
| Ne categorization | Number in category | Percent of total | Metric mean value |
| Low | 12 | 5.26 | -5.05 |
| Medium | 38 | 16.67 | -3.69 |
| High | 178 | 78.07 | -2.45 |
| Total | 228 | 100 | -2.80 |

**Table 3.** SpecKS parameters used in dataset 1.

| **Table 3. SpecKS parameters used in dataset 1** |  |  |
| --- | --- | --- |
| Polyploid | DIV_time_MYA | [80,70, 60, 50, 40, 30, 20,10] |
| Polyploid | WGD_time_MYA | [0,5,10,20,50] |
| Polyploid | gene_div_time_distribution_parameters | "Impulse,1,1" and "expo,0,K"<br>Where K varied from<br>[0.01,0.1,1.0,5.0,10.0,20.0]<br>To span [Gossmann 2010] |
| Species Tree | full_sim_time | 100 MY |
| Gene Tree | mean_gene_birth_rate_GpMY | 0.001359<br>genes per MY |
| Gene Tree | SSD_half_life_MY | 4 MY |
| Gene Tree | WGD_half_life_MY | 31 MY |
| Gene Tree | num_gene_trees_per_species_tree | 3000 |
| Sequence Evolution | num_replicates_per_gene_tree | 1 |
| Sequence Evolution | num_codons | 1000 |
| Sequence Evolution | Ks_per_Myr | 0.01 |
| Sequence Evolution | per_site_evolutionary_distance | 0.01268182 |
| All other parameters are default. |  |  |

Running SpecKS: SpecKS is run calling the “SpecKS.py” function from python and passing it the config xml file. For example, “python3 SpecKS.py myconfig.xml”

SpecKS output: The SpecKS output file is a .csv file, with each line giving information for a unique gene-pair. The data are organized into 4 columns, giving the gene-tree name of the pair, the names of the two genes in the pair (the leaves of the tree), the *Ks* value, and the path to the CODEML output file from which the value originated,respectively.

Parallelization: For simplicity, SpecKS is not internally parallelized. A single polyploid of about 3000 genes, 1000 codons long, simulated over 100 MY, runs on a typical cluster (e.g. 16-node Dual-10 Xeon CPU, E5 -2630v4 2.20GHz, 12.8GB RAM per core) in under 10 minutes. To simulate batches of polyploids, we recommend submitting each polyploid as a separate batch job to a job scheduler such as SGE (Oracle) or SLURM (SchedMD).

## Source code and data availability

SpecKS is available from the github repository https://github.com/tamsen/SpecKS. SpecKS input files used to drive the simulation and the scripts used to generate the figures are at https://github.com/tamsen/SpecKS_paper_scripts.

## Acknowledgments

We would like to thank members of the Barker and Sethuraman Labs for valuable inputs during the conceptualization stage of this project. We would like to acknowledge Matthew Hahn and two anonymous reviewers for valuable suggestions on the first version of this manuscript. AS and TD were both supported by NSF CAREER: 2147812 and two CSUBIOTECH grants. All computations were performed on the *mesxuuyan* cluster at San Diego State University, which was supported by startup funds and NSF ABI: 1564659 to AS.

## References

Al-Dossary O, Furtado A, KharabianMasouleh A, Alsubaie B, Al-Mssallem I, Henry RJ. 2023. Long read sequencing to reveal the full complexity of a plant transcriptome by targeting both standard and long workflows. Plant Methods 19:112.

Arrigo N, Barker MS. 2012. Rarely successful polyploids and their legacy in plant genomes. Current Opinion in Plant Biology 15:140–146.

Baduel P, Bray S, Vallejo-Marin M, Kolář F, Yant L. 2018. The “polyploid hop”: shifting challenges and opportunities over the evolutionary lifespan of genome duplications. Frontiers in Ecology and Evolution 6:117.

Baniaga AE, Marx HE, Arrigo N, Barker MS. 2020. Polyploid plants have faster rates of multivariate niche differentiation than their diploid relatives. Ecology Letters 23:68–78.

Barba-Montoya J, Dos Reis M, Schneider H, Donoghue PCJ, Yang Z. 2018. Constraining uncertainty in the timescale of angiosperm evolution and the veracity of a Cretaceous Terrestrial Revolution. New Phytol 218:819–834.

Barker MS, Arrigo N, Baniaga AE, Li Z, Levin DA. 2016. On the relative abundance of autopolyploids and allopolyploids. New Phytologist 210:391–398.

Blanc G, Wolfe KH. 2004. Widespread paleopolyploidy in model plant species inferred from age distributions of duplicate genes. Plant Cell 16:1667.

Blischak PD, Sajan M, Barker MS, Gutenkunst RN. 2023. Demographic history inference and the polyploid continuum. Genetics 224:iyad107.

Bomblies K, Jones G, Franklin C, Zickler D, Kleckner N. 2016. The challenge of evolving stable polyploidy: could an increase in “crossover interference distance” play a central role? Chromosoma 125:287–300.

Booker WW, Schrider DR. 2023. The genetic consequences of range expansion and its influence on diploidization in polyploids. bioRxiv [Internet]. Available from: https://www.biorxiv.org/content/early/2023/10/21/2023.10.18.562992

Bouckaert R, Vaughan TG, Barido-Sottani J, Duchêne S, Fourment M, Gavryushkina A, Heled J, Jones G, Kühnert D, De Maio N, et al. 2019. BEAST 2.5: An advanced software platform for Bayesian evolutionary analysis. PLoS Comput Biol 15:e1006650.

Bowers JE, Chapman BA, Rong J, Paterson AH. 2003. Unravelling angiosperm genome evolution by phylogenetic analysis of chromosomal duplication events. Nature 422:433– 438.

Buggs RJA, Zhang L, Miles N, Tate JA, Gao L, Wei W, Schnable PS, Barbazuk WB, Soltis PS, Soltis DE. 2011. Transcriptomic shock generates evolutionary novelty in a newly formed, natural allopolyploid plant. Curr Biol 21:551–556.

Carretero-Paulet L, Van de Peer Y. 2020. The evolutionary conundrum of whole-genome duplication. Am J Bot 107:1101–1105.

Chen H, Zwaenepoel A. 2023. Inference of ancient polyploidy from genomic data. Methods Mol Biol 2545:3–18.

Clark JW, Donoghue PC. 2018. Whole-genome duplication and plant macroevolution. Trends in Plant Science 23:933–945.

Clark JW, Donoghue PC. 2023. Constraining whole-genome duplication events in geological time. In: Polyploidy: Methods and Protocols. Springer. p. 139–154.

Clark JW, Puttick MN, Donoghue PCJ. 2019. Origin of horsetails and the role of whole-genome duplication in plant macroevolution. Proc Biol Sci 286:20191662.

Clausen J, Keck DD, Hiesey WM. 1945. Experimental studies on the nature of species. 2. Plant evolution through amphiploidy and autoploidy, with examples from the Madiinae. Experimental studies on the nature of species. 2. Plant evolution through amphiploidy and autoploidy, with examples from the Madiinae.

Comai L, Tyagi AP, Winter K, Holmes-Davis R, Reynolds SH, Stevens Y, Byers B. 2000. Phenotypic instability and rapid gene silencing in newly formed arabidopsis allotetraploids. Plant Cell 12:1551.

Conant GC. 2023. POInT: Modeling polyploidy in the era of ubiquitous genomics. Methods Mol Biol 2545:77–90.

Cui L, Wall PK, Leebens-Mack JH, Lindsay BG, Soltis DE, Doyle JJ, Soltis PS, Carlson JE, Arumuganathan K, Barakat A, et al. 2006. Widespread genome duplications throughout the history of flowering plants. Genome Res 16:738–749.

Dai X, Hu Q, Cai Q, Feng K, Ye N, Tuskan GA, Milne R, Chen Y, Wan Z, Wang Z, et al. 2014. The willow genome and divergent evolution from poplar after the common genome duplication. Cell Research 24:1274–1277.

Darlington CD. 1937. Recent advances in cytology. Recent advances in cytology.

David KT, Oaks JR, Halanych KM. 2020. Patterns of gene evolution following duplications and speciations in vertebrates. PeerJ 8:e8813.

De La Torre AR, Li Z, Van de Peer Y, Ingvarsson PK. 2017. Contrasting rates of molecular evolution and patterns of selection among gymnosperms and flowering plants. Molecular Biology and Evolution 34:1363–1377.

Deb SK, Edger PP, Pires JC, McKain MR. 2023. Patterns, mechanisms, and consequences of homoeologous exchange in allopolyploid angiosperms: a genomic and epigenomic perspective. New Phytologist.

Dodsworth S, Chase MW, Leitch AR. 2016. Is post-polyploidization diploidization the key to the evolutionary success of angiosperms? Botanical Journal of the Linnean Society 180:1–5.

Douglas GM, Gos G, Steige KA, Salcedo A, Holm K, Josephs EB, Arunkumar R, Ågren JA, Hazzouri KM, Wang W. 2015. Hybrid origins and the earliest stages of diploidization in the highly successful recent polyploid *Capsella bursa-pastoris*. Proceedings of the National Academy of Sciences 112:2806–2811.

Doyle JJ, Egan AN. 2010. Dating the origins of polyploidy events. New Phytol 186:73–85.

Doyle JJ, Sherman-Broyles S. 2017. Double trouble: taxonomy and definitions of polyploidy. New Phytologist 213:487–493.

Estep MC, McKain MR, Diaz DV, Zhong J, Hodge JG, Hodkinson TR, Layton DJ, Malcomber ST, Pasquet R, Kellogg EA. 2014. Allopolyploidy, diversification, and the Miocene grassland expansion. Proceedings of the National Academy of Sciences 111:15149– 15154.

Gaeta RT, Chris Pires J. 2010. Homoeologous recombination in allopolyploids: the polyploid ratchet. New Phytologist 186:18–28.

Garsmeur O, Schnable JC, Almeida A, Jourda C, D’Hont A, Freeling M. 2014. Two Evolutionarily Distinct Classes of Paleopolyploidy. Molecular Biology and Evolution 31:448–454.

Gaut BS, Doebley JF. 1997. DNA sequence evidence for the segmental allotetraploid origin of maize. Proceedings of the National Academy of Sciences 94:6809–6814.

Gaut BS, Seymour DK, Liu Q, Zhou Y. 2018. Demography and its effects on genomic variation in crop domestication. Nature Plants 4:512–520.

Goldman N, Yang Z. 1994. A codon-based model of nucleotide substitution for protein-coding DNA sequences. Mol Biol Evol 11:725–736.

Gossmann TI, Song B-H, Windsor AJ, Mitchell-Olds T, Dixon CJ, Kapralov MV, Filatov DA, Eyre-Walker A. 2010. Genome wide analyses reveal little evidence for adaptive evolution in many plant species. Mol Biol Evol 27:1822–1832.

Guo X, Mandáková T, Trachtová K, Özüdoğru B, Liu J, Lysak MA. 2020. Linked by ancestral bonds: Multiple whole-genome duplications and reticulate evolution in a Brassicaceae tribe. Molecular Biology and Evolution 38:1695–1714.

Guo Y-L. 2013. Gene family evolution in green plants with emphasis on the origination and evolution of *Arabidopsis thaliana* genes. Plant J 73:941–951.

Gutenkunst RN, Hernandez RD, Williamson SH, Bustamante CD. 2009. Inferring the joint demographic history of multiple populations from multidimensional SNP frequency data. PLoS genetics 5:e1000695.

Hampson S, McLysaght A, Gaut B, Baldi P. 2003. LineUp: statistical detection of chromosomal homology with application to plant comparative genomics. Genome Res 13:999–1010.

Hoegg S, Brinkmann H, Taylor JS, Meyer A. 2004. Phylogenetic timing of the fish-specific genome duplication correlates with the diversification of teleost fish. Journal of Molecular Evolution 59:190–203.

Hunter JD. 2007. Matplotlib: A 2D graphics environment. Computing in science & engineering 9:90–95.

Kingman JFC. 1982. The coalescent. Stochastic Processes and their Applications 13:235–248.

Lallemand T, Leduc M, Desmazières A, Aubourg S, Rizzon C, Landès C, Celton J-M. 2023. Insights into the evolution of ohnologous sequences and their epigenetic marks post-WGD in *Malus domestica*. Genome Biology and Evolution 15:evad178.

Laurent S, Salamin N, Robinson-Rechavi M. 2017. No evidence for the radiation time lag model after whole genome duplications in Teleostei. PLoS One 12:e0176384–e0176384.

Le Comber SC, Ainouche ML, Kovarik A, Leitch AR. 2010. Making a functional diploid: from polysomic to disomic inheritance. New Phytol 186:113–122.

Leebens-Mack JH, Barker MS, Carpenter EJ, Deyholos MK, Gitzendanner MA, Graham SW, Grosse I, Li Z, Melkonian M, Mirarab S, et al. 2019. One thousand plant transcriptomes and the phylogenomics of green plants. Nature 574:679–685.

Levin DA. 2013. The timetable for allopolyploidy in flowering plants. Ann Bot 112:1201–1208.

Li Z, Baniaga AE, Sessa EB, Scascitelli M, Graham SW, Rieseberg LH, Barker MS. 2015. Early genome duplications in conifers and other seed plants. Science advances 1:e1501084.

Li Z, Barker MS. 2020. Inferring putative ancient whole-genome duplications in the 1000 Plants (1KP) initiative: access to gene family phylogenies and age distributions. GigaScience 9:giaa004.

Li Z, McKibben MTW, Finch GS, Blischak PD, Sutherland BL, Barker MS. 2021. Patterns and Processes of Diploidization in Land Plants. Annual Review of Plant Biology 72:387–410.

Li Z, Tiley GP, Galuska SR, Reardon CR, Kidder TI, Rundell RJ, Barker MS. 2018. Multiple large-scale gene and genome duplications during the evolution of hexapods. Proceedings of the National Academy of Sciences 115:4713–4718.

Lott M, Spillner A, Huber KT, Moulton V. 2009. PADRE: a package for analyzing and displaying reticulate evolution. Bioinformatics 25:1199–1200.

Luo D, Zhou Q, Wu Y, Chai X, Liu W, Wang Y, Yang Q, Wang Z, Liu Z. 2019. Full-length transcript sequencing and comparative transcriptomic analysis to evaluate the contribution of osmotic and ionic stress components towards salinity tolerance in the roots of cultivated alfalfa (Medicago sativa L.). BMC Plant Biology 19:32.

Lv Z, Addo Nyarko C, Ramtekey V, Behn H, Mason AS. 2024. Defining autopolyploidy: Cytology, genetics, and taxonomy. American Journal of Botany:e16292.

Lynch M, Conery JS. 2000. The evolutionary fate and consequences of duplicate genes. Science 290:1151–1155.

Lynch M, Conery JS. 2003. The evolutionary demography of duplicate genes. Genome Evolution: Gene and Genome Duplications and the Origin of Novel Gene Functions:35– 44.

Lynch M, O’Hely M, Walsh B, Force A. 2001. The probability of preservation of a newly arisen gene duplicate. Genetics 159:1789–1804.

Mable BK. 2013. Polyploids and hybrids in changing environments: winners or losers in the struggle for adaptation? Heredity 110:95–96.

Madlung A. 2013. Polyploidy and its effect on evolutionary success: old questions revisited with new tools. Heredity 110:99–104.

Maere S, De Bodt S, Raes J, Casneuf T, Van Montagu M, Kuiper M, Van de Peer Y. 2005. Modeling gene and genome duplications in eukaryotes. Proc Natl Acad Sci U S A 102:5454–5459.

Mason AS, Wendel JF. 2020. Homoeologous exchanges, segmental allopolyploidy, and polyploid genome evolution. Frontiers in Genetics:1014.

Mayer VW, Aguilera A. 1990. High levels of chromosome instability in polyploids of Saccharomyces cerevisiae. Mutation Research/Fundamental and Molecular Mechanisms of Mutagenesis 231:177–186.

Mayfield-Jones D, Washburn JD, Arias T, Edger PP, Pires JC, Conant GC. 2013. Watching the grin fade: tracing the effects of polyploidy on different evolutionary time scales. Semin Cell Dev Biol 24:320–331.

Mccann J, Jang T-S, Macas J, Schneeweiss GM, Matzke NJ, Novák P, Stuessy TF, Villaseñor JL, Weiss-Schneeweiss H. 2018. Dating the species network: Allopolyploidy and repetitive DNA evolution in American daisies (*Melampodium sect. Melampodium*, Asteraceae). Syst Biol 67:1010–1024.

McClintock B. 1929. A cytological and genetical study of triploid maize. Genetics 14:180.

Meirmans P, Van Tienderen P. 2013. The effects of inheritance in tetraploids on genetic diversity and population divergence. Heredity 110:131–137.

Nieto Feliner G, Casacuberta J, Wendel JF. 2020. Genomics of evolutionary novelty in hybrids and polyploids. Frontiers in Genetics 11:792.

Ohno S. 1970. Evolution by gene duplication. Berlin, New York: Berlin, New York, Springer-Verlag

Otto SP, Whitton J. 2000. Polyploid incidence and evolution. Annu. Rev. Genet. 34:401–437.

Parey E, Louis A, Cabau C, Guiguen Y, Roest Crollius H, Berthelot C. 2020. Synteny-Guided Resolution of Gene Trees Clarifies the Functional Impact of Whole-Genome Duplications. Mol Biol Evol 37:3324–3337.

Parey E, Louis A, Montfort J, Guiguen Y, Crollius HR, Berthelot C. 2022. An atlas of fish genome evolution reveals delayed rediploidization following the teleost whole-genome duplication. Genome Research 32:1685–1697.

Parisod C, Holderegger R, Brochmann C. 2010. Evolutionary consequences of autopolyploidy. New Phytologist 186:5–17.

Pedregosa F, Varoquaux G, Gramfort A, Michel V, Thirion B, Grisel O, Blondel M, Prettenhofer P, Weiss R, Dubourg V. 2011. Scikit-learn: Machine learning in Python. the Journal of machine Learning research 12:2825–2830.

Qiao X, Li Q, Yin H, Qi K, Li L, Wang R, Zhang S, Paterson AH. 2019. Gene duplication and evolution in recurring polyploidization–diploidization cycles in plants. Genome Biology 20:38.

Ramsey J, Schemske DW. 2002. Neopolyploidy in flowering plants. Annu. Rev. Ecol. Syst. 33:589–639.

Ren R, Wang H, Guo C, Zhang N, Zeng L, Chen Y, Ma H, Qi J. 2018. Widespread whole genome duplications contribute to genome complexity and species diversity in angiosperms. Molecular Plant 11:414–428.

Robertson FM, Gundappa MK, Grammes F, Hvidsten TR, Redmond AK, Lien S, Martin SAM, Holland PWH, Sandve SR, Macqueen DJ. 2017. Lineage-specific rediploidization is a mechanism to explain time-lags between genome duplication and evolutionary diversification. Genome Biology 18:111.

Rothfels CJ, Johnson AK, Hovenkamp PH, Swofford DL, Roskam HC, Fraser-Jenkins CR, Windham MD, Pryer KM. 2015. Natural hybridization between genera that diverged from each other approximately 60 million years ago. Am Nat 185:433–442.

Roux C, Pannell JR. 2015. Inferring the mode of origin of polyploid species from next-generation sequence data. Mol Ecol 24:1047–1059.

Roux J, Liu J, Robinson-Rechavi M. 2017. Selective constraints on coding sequences of nervous system genes are a major determinant of duplicate gene retention in vertebrates. Molecular biology and evolution 34:2773–2791.

Salojarvi J, Rambani A, Yu Z, Guyot R, Strickler S, Lepelley M, Wang C, Rajaraman S, Rastas P, Zheng C. 2023. The genome and population genomics of allopolyploid *Coffea arabica* reveal the diversification history of modern coffee cultivars. bioRxiv:2023.09. 06.556570.

Salojärvi J, Rambani A, Yu Z, Guyot R, Strickler S, Lepelley M, Wang C, Rajaraman S, Rastas P, Zheng C. 2024. The genome and population genomics of allopolyploid Coffea arabica reveal the diversification history of modern coffee cultivars. Nature genetics 56:721–731.

Schlueter JA, Dixon P, Granger C, Grant D, Clark L, Doyle JJ, Shoemaker RC. 2004. Mining EST databases to resolve evolutionary events in major crop species. Genome 47:868– 876.

Schranz E, Mohammadin S, Edger PP. 2012. Ancient whole genome duplications, novelty and diversification: the WGD Radiation Lag-Time Model. Current Opinion in Plant Biology 15:147–153.

Senchina DS, Alvarez I, Cronn RC, Liu B, Rong J, Noyes RD, Paterson AH, Wing RA, Wilkins TA, Wendel JF. 2003. Rate variation among nuclear genes and the age of polyploidy in Gossypium. Mol Biol Evol 20:633–643.

Sensalari C, Maere S, Lohaus R. 2022. ksrates: positioning whole-genome duplications relative to speciation events in KS distributions. Bioinformatics 38:530–532.

Soltis DE, Buggs RJA, Doyle JJ, Soltis PS. 2010. What we still don’t know about polyploidy. TAXON 59:1387–1403.

Soltis DE, Soltis PS, Schemske DW, Hancock JF, Thompson JN, Husband BC, Judd WS. 2007. Autopolyploidy in angiosperms: have we grossly underestimated the number of species? Taxon 56:13–30.

Soltis DE, Visger CJ, Marchant DB, Soltis PS. 2016. Polyploidy: pitfalls and paths to a paradigm. American Journal of Botany 103:1146–1166.

Soltis PS, Liu X, Marchant DB, Visger CJ, Soltis DE. 2014. Polyploidy and novelty: Gottlieb’s legacy. Philos Trans R Soc Lond B Biol Sci 369:20130351.

Soltis PS, Soltis DE. 2012. Polyploidy and genome evolution. Springer

Spoelhof JP, Soltis PS, Soltis DE. 2017. Pure polyploidy: closing the gaps in autopolyploid research. Journal of Systematics and Evolution 55:340–352.

St Onge KR, Foxe JP, Li J, Li H, Holm K, Corcoran P, Slotte T, Lascoux M, Wright SI. 2012. Coalescent-based analysis distinguishes between allo- and autopolyploid origin in Shepherd’s Purse (*Capsella bursa-pastoris*). Mol Biol Evol 29:1721–1733.

Stebbins CL. 1950. Variation and evolution in plants. *Variation and evolution in plants*.

Stebbins GL. 1947. Types of polyploids: their classification and significance. In: Advances in genetics. Vol. 1. Elsevier. p. 403–429.

Stebbins GL. 1951. Cataclysmic Evolution. Scientific American 184:54–59.

Sutherland BL, Tiley GP, Li Z, McKibben MT, Barker MS. 2024. SLEDGe: Inference of ancient whole genome duplications using machine learning. bioRxiv:2024.01.17.574559.

Tank DC, Eastman JM, Pennell MW, Soltis PS, Soltis DE, Hinchliff CE, Brown JW, Sessa EB, Harmon LJ. 2015. Nested radiations and the pulse of angiosperm diversification: increased diversification rates often follow whole genome duplications. New Phytologist 207:454–467.

Thomas GWC, Ather SH, Hahn MW. 2017. Gene-tree reconciliation with MUL-trees to resolve polyploidy events. Systematic Biology 66:1007–1018.

Tiley GP, Barker MS, Burleigh JG. 2018. Assessing the performance of Ks plots for detecting ancient whole genome duplications. Genome Biology and Evolution 10:2882–2898.

Van de Peer Y. 2004. Computational approaches to unveiling ancient genome duplications. Nature Reviews Genetics 5:752–763.

Van de Peer Y, Ashman T-L, Soltis PS, Soltis DE. 2020. Polyploidy: an evolutionary and ecological force in stressful times. The Plant Cell [Internet]. Available from: 10.1093/plcell/koaa015

Vandepoele K, Saeys Y, Simillion C, Raes J, Van De Peer Y. 2002. The automatic detection of homologous regions (ADHoRe) and its application to microcolinearity between Arabidopsis and rice. Genome Res 12:1792–1801.

Vanneste K, Van de Peer Y, Maere S. 2013. Inference of genome duplications from age distributions revisited. Mol Biol Evol 30:177–190.

Wang J, Qin J, Sun P, Ma X, Yu J, Li Y, Sun S, Lei T, Meng F, Wei C. 2019. Polyploidy index and its implications for the evolution of polyploids. Frontiers in genetics 10:807.

Wang X, Shi X, Li Z, Zhu Q, Kong L, Tang W, Ge S, Luo J. 2006. Statistical inference of chromosomal homology based on gene colinearity and applications to Arabidopsis and rice. BMC Bioinformatics 7:447.

Wen D, Yu Y, Zhu J, Nakhleh L. 2018. Inferring phylogenetic networks using phyloNet. Systematic Biology 67:735–740.

Wendel JF. 2015. The wondrous cycles of polyploidy in plants. Am J Bot 102:1753–1756.

Wendel JF, Jackson SA, Meyers BC, Wing RA. 2016. Evolution of plant genome architecture. Genome Biology 17:37.

Wolfe KH, Li WH, Sharp PM. 1987. Rates of nucleotide substitution vary greatly among plant mitochondrial, chloroplast, and nuclear DNAs. Proc Natl Acad Sci U S A 84:9054–9058.

Yan Z, Cao Z, Liu Y, Ogilvie HA, Nakhleh L. 2022. Maximum parsimony inference of phylogenetic networks in the presence of polyploid complexes. Systematic Biology 71:706–720.

Yang N, Wang Y, Liu X, Jin M, Vallebueno-Estrada M, Calfee E, Chen L, Dilkes BP, Gui S, Fan X, et al. 2023. Two teosintes made modern maize. Science 382:eadg8940.

Yang Z. 2007. PAML 4: phylogenetic analysis by maximum likelihood. Molecular Biology and Evolution 24:1586–1591.

Yang Z, Nielsen R. 1998. Synonymous and nonsynonymous rate variation in nuclear genes of mammals. Journal of Molecular Evolution 46:409–418.

Yang Z, Nielsen R. 2000. Estimating synonymous and nonsynonymous substitution rates under realistic evolutionary models. Mol Biol Evol 17:32–43.

Yant L, Bomblies K. 2015. Genome management and mismanagement--cell-level opportunities and challenges of whole-genome duplication. Genes Dev 29:2405–2419.

Yu Q, Guyot R, de Kochko A, Byers A, Navajas-Pérez R, Langston BJ, Dubreuil-Tranchant C, Paterson AH, Poncet V, Nagai C, et al. 2011. Micro-collinearity and genome evolution in the vicinity of an ethylene receptor gene of cultivated diploid and allotetraploid coffee species (Coffea). The Plant Journal 67:305–317.

Zhang H, Gou X, Zhang A, Wang X, Zhao N, Dong Y, Li L, Liu B. 2016. Transcriptome shock invokes disruption of parental expression-conserved genes in tetraploid wheat. Scientific Reports 6:26363.

Zhang L, Wu S, Chang X, Wang X, Zhao Y, Xia Y, Trigiano RN, Jiao Y, Chen F. 2020. The ancient wave of polyploidization events in flowering plants and their facilitated adaptation to environmental stress. *Plant*, Cell & Environment 43:2847–2856.

Zhang S, Li R, Zhang L, Chen S, Xie M, Yang L, Xia Y, Foyer CH, Zhao Z, Lam H-M. 2020. New insights into *Arabidopsis* transcriptome complexity revealed by direct sequencing of native RNAs. Nucleic Acids Res 48:7700–7711.

Zhao J, Teufel AI, Liberles DA, Liu L. 2015. A generalized birth and death process for modeling the fates of gene duplication. BMC Evol Biol 15:275.

Zwaenepoel A, Van de Peer Y. 2019. WGD—simple command line tools for the analysis of ancient whole-genome duplications. Bioinformatics 35:2153–2155.

