## Supplementary Figures and Tables for "Accurate Inference of the Polyploid Continuum using Forward-time Simulations"

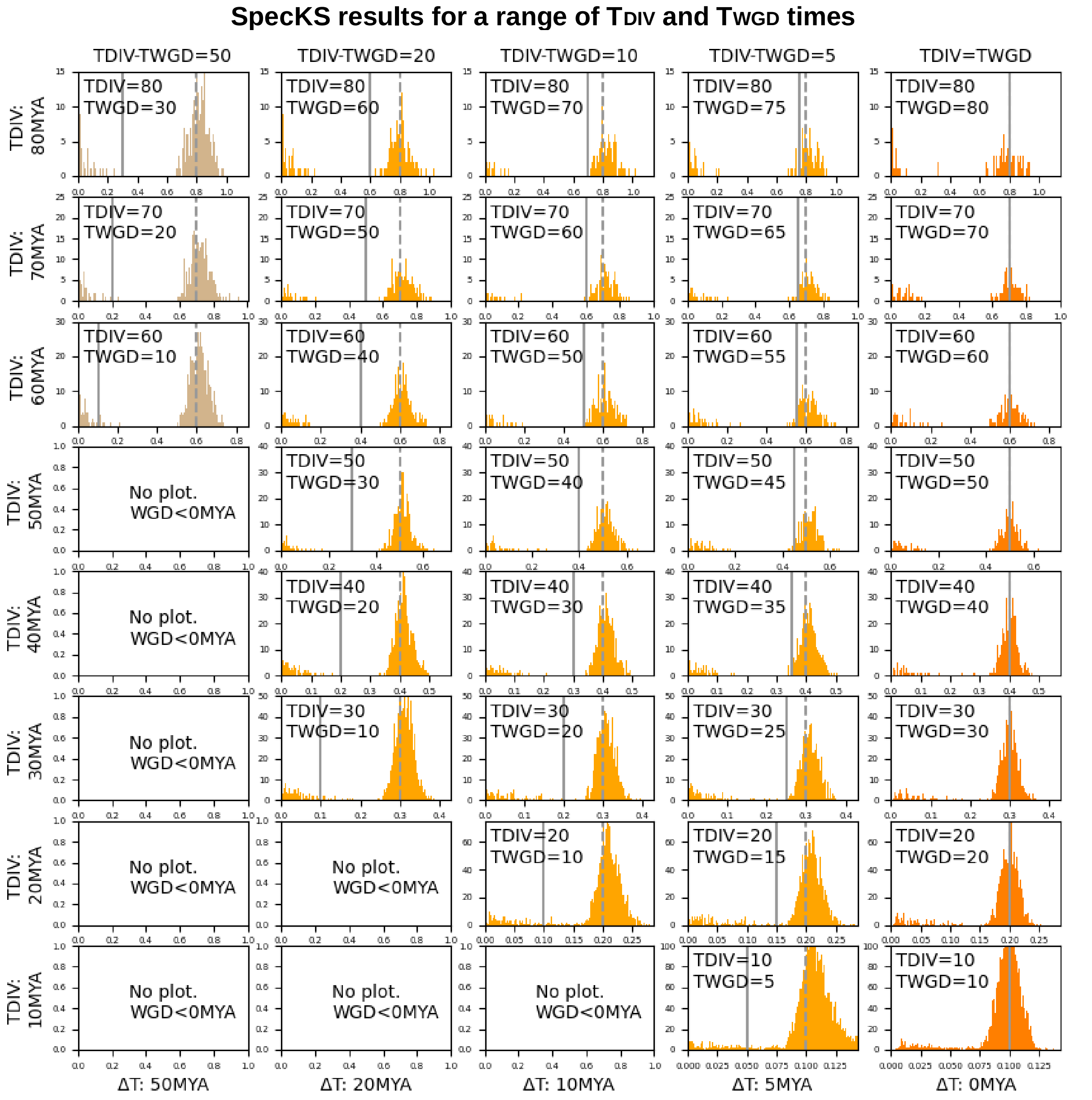


**Figure S1. SpecKS gives realistic Ks histograms for a range of TDIV and TWGD times.** TDIV times range from 10-80 MYA (the rows), and TWGD is offset from speciation time by 50, 20,10, 5 and 0 MY (the columns). Autopolyploids are the right-most column, in red, and allopolyploids are blue. The x-axis is *Ks* and the y-axis is the number of paralogs. The *Ks*-equivalent TWGD and TDIV times are given in gray, with WDG being the solid line, and DIV time dashed. $N_e$*$G_t$ was held fixed at 1MY for ancestral allopolyploid *Ks* distributions.

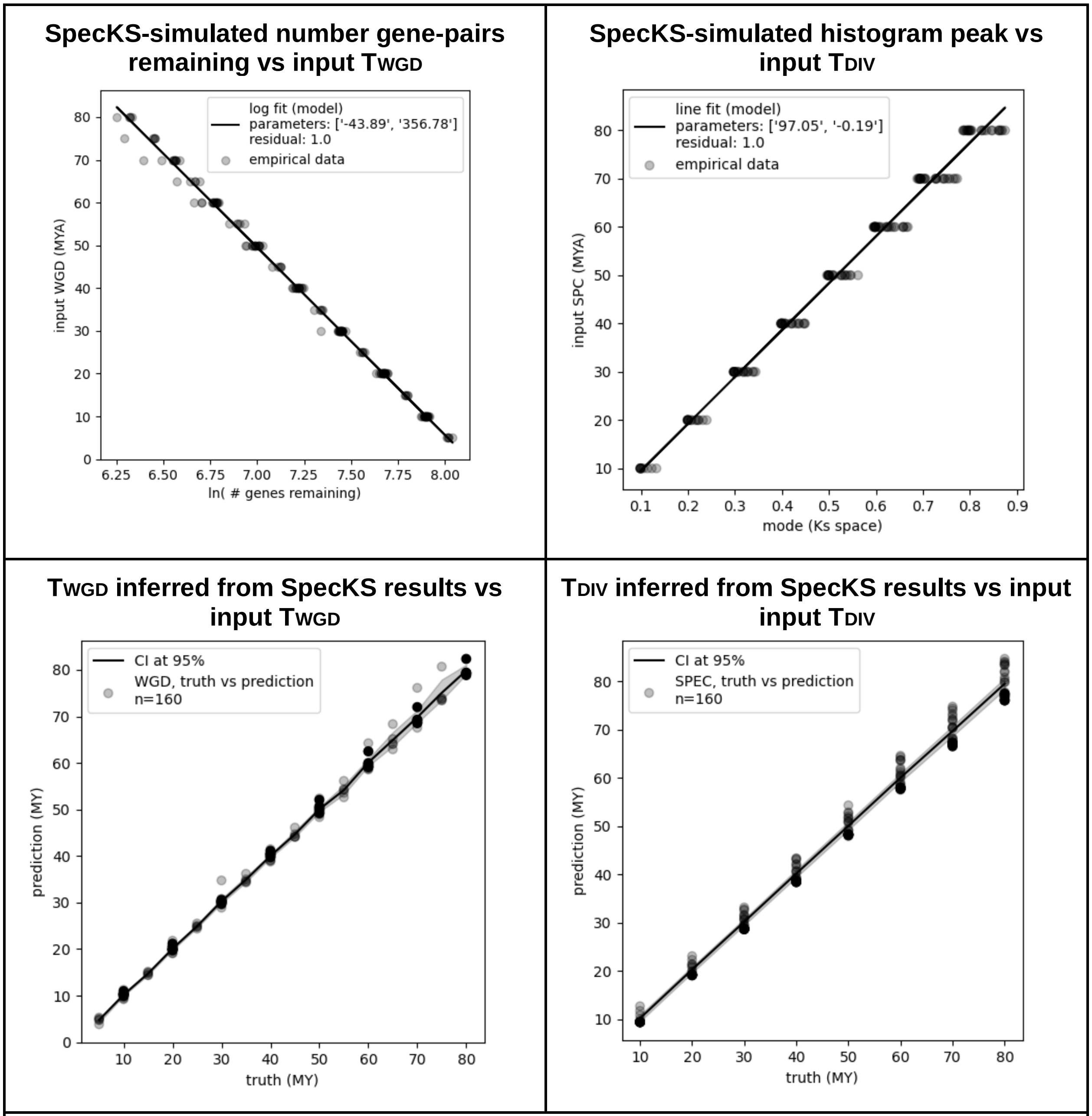


**Figure S2. SpecKS allows for the accurate recovery of the TWGD and TDIV.**
Top left: SpecKS-simulated histogram mode (x-axis) vs input TWGD time (y-axis). Top right: SpecKS-simulated natural log of the number of gene pairs remaining (x-axis) vs input TWGD time (y-axis). Bottom left: TDIV inferred from SpecKS results (y-axis), vs input TDIV (x-axis). Bottom right: TWGD time inferred from SpecKS results (y-axis), vs input TWGD (x-axis). Each dot represents results from a simulated polyploid histogram.

**Histograms from the 1KP dataset, as categorized by our high-vs-low Ne discriminator**

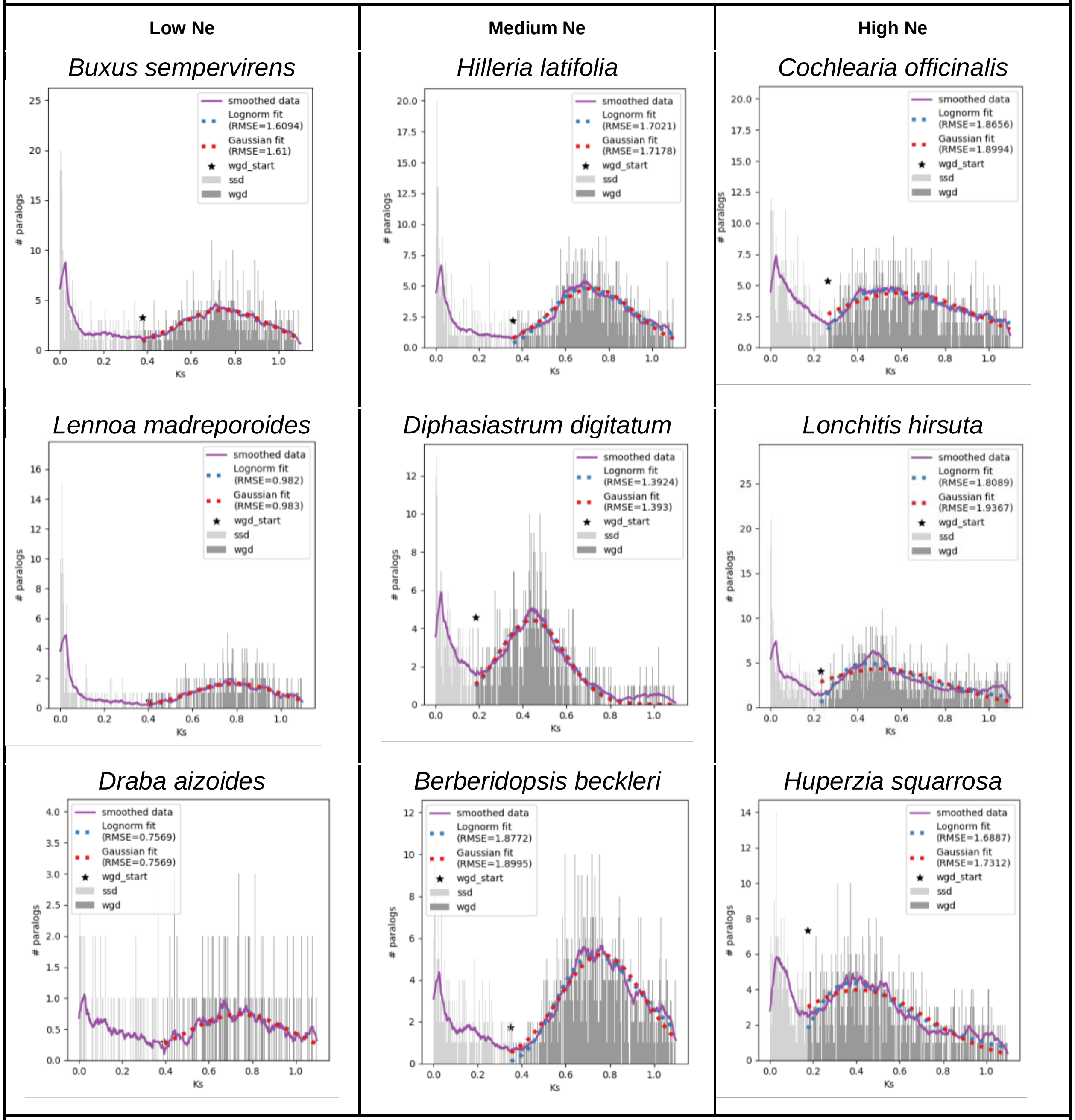


**Figure S3. *Ks* histograms for example species from the 1KP dataset, as categorized by the high-vs-low Ne discrimination model**. X-axis is *Ks*. Y-axis is the number of paralog pairs. We show that the *Ks* distributions of species categorized as low-Ne have more more symmetric *Ks* distributions (the best lognormal fit is identically the Gaussian) as opposed to the high-Ne samples (where a fat-tailed lognormal is a better fit than the Gaussian).

Supplementary Tables

| **Table S1: All categorization results from the high-vs-low Ne discrimination model to *Ks* histograms from the 1KP dataset.** | | |
|---|---|---|
| **Species** | **Classification** | **Score** |
| *Draba aizoides* | Low | -6.546697946 |
| *Buxus sempervirens* | Low | -5.717641879 |
| *Saruma henryi* | Low | -5.565407938 |
| *Sabal bermudana* | Low | -5.237591203 |
| *Lennoa madreporoides* | Low | -5.229114357 |
| *Astragalus membranaceus* | Low | -4.885065177 |
| *Sarcandra glabra* | Low | -4.858459511 |
| *Anticharis glandulosa* | Low | -4.744451134 |
| *Pseudotaxiphyllum elegans* | Low | -4.511522573 |
| *Acacia argyrophylla* | Low | -4.489601599 |
| *Hedwigia ciliata* | Low | -4.465070208 |
| *Helenium autumnale* | Low | -4.39941065 |
| *Ipomoea purpurea* | Medium | -4.273574798 |
| *Ipomoea indica* | Medium | -4.264493558 |
| *Talinum sp.* | Medium | -4.239670051 |
| *Lycopodium deuterodensum* | Medium | -4.10780315 |
| *Anomodon attenuatus* | Medium | -3.973239661 |
| *Conopholis americana* | Medium | -3.938689476 |
| *Cercidiphyllum japonicum* | Medium | -3.913510393 |
| *Convolvulus arvensis* | Medium | -3.866300875 |

| | | |
|---|---|---|
| *Blutaparon vermiculare* | Medium | -3.849987869 |
| *Saponaria officinalis* | Medium | -3.829760148 |
| *Oxalis sp.* | Medium | -3.82198443 |
| *Aextoxicon punctatum* | Medium | -3.804071819 |
| *Lomandra longifolia* | Medium | -3.801791645 |
| *Diphasiastrum digitatum* | Medium | -3.748671419 |
| *Phacelia campanularia* | Medium | -3.734868012 |
| *Linum grandiflorum* | Medium | -3.706705824 |
| *Lycopodium annotinum* | Medium | -3.68907275 |
| *Solanum xanthocarpum* | Medium | -3.666454134 |
| *Lonicera japonica* | Medium | -3.665831976 |
| *Typha latifolia* | Medium | -3.655680917 |
| *Polygonum convolvulus* | Medium | -3.635038174 |
| *Mansoa alliacea* | Medium | -3.625956664 |
| *Strychnos spinosa* | Medium | -3.586528125 |
| *Ipomoea nil* | Medium | -3.55770231 |
| *Typha angustifolia* | Medium | -3.556129656 |
| *Linum hirsutum* | Medium | -3.52201674 |
| *Berberidopsis beckleri* | Medium | -3.50122304 |
| *Ascarina rubricaulis* | Medium | -3.480674241 |
| *Begonia sp.* | Medium | -3.473109533 |
| *Symphoricarpos sp.* | Medium | -3.468799615 |

| | | |
|---|---|---|
| *Tabebuia umbellata* | Medium | -3.461661429 |
| *Ipomoea coccinea* | Medium | -3.446163284 |
| *Ipomoea lobata* | Medium | -3.436296906 |
| *Dendrolycopodium obscurum* | Medium | -3.423820489 |
| *Pholisma arenarium* | Medium | -3.417761648 |
| *Ipomoea hederacea* | Medium | -3.400176853 |
| *Hilleria latifolia* | Medium | -3.392231237 |
| *Antirrhinum braun-blanquetii* | Medium | -3.387758654 |
| *Solanum lasiophyllum* | High | -3.337128128 |
| *Codariocalyx motorius* | High | -3.323662392 |
| *Silene latifolia* | High | -3.299954965 |
| *Ledum palustre* | High | -3.283271441 |
| *Centella asiatica* | High | -3.264483834 |
| *Lindsaea linearis* | High | -3.251731762 |
| *Pseudolycopodiella caroliniana* | High | -3.236745736 |
| *Racomitrium elongatum* | High | -3.228940525 |
| *Salix fargesii* | High | -3.214291889 |
| *Vitex agnus-castus* | High | -3.207670647 |
| *Ilex paraguariensis* | High | -3.202042481 |
| *Tropaeolum peregrinum* | High | -3.179356871 |
| *Passiflora caerulea* | High | -3.163989265 |
| *Bauhinia tomentosa* | High | -3.149160182 |

| | | |
|---|---|---|
| *Leucobryum albidum* | High | -3.142131876 |
| *Fouquieria macdougalii* | High | -3.115979346 |
| *Galax urceolata* | High | -3.101156547 |
| *Diospyros malabarica* | High | -3.097618909 |
| *Manihot grahamii* | High | -3.075376302 |
| *Erythroxylum coca* | High | -3.072872827 |
| *Ipomoea quamoclit* | High | -3.071285007 |
| *Salix purpurea* | High | -3.06507262 |
| *Rhizophora mangle* | High | -3.053472345 |
| *Paulownia fargesii* | High | -3.05179418 |
| *Petiveria alliacea* | High | -3.049704367 |
| *Sinningia tuberosa* | High | -3.046477226 |
| *Galax urceolata* | High | -3.023315417 |
| *Idiospermum australiense* | High | -3.022708763 |
| *Salix viminalis* | High | -2.995387547 |
| *Dendropemon caribaeus* | High | -2.976001421 |
| *Drypetes deplanchei* | High | -2.974300131 |
| *Uncarina grandidieri* | High | -2.97406581 |
| *Lycium sp.* | High | -2.959174934 |
| *Ehretia acuminata* | High | -2.956496422 |
| *Lycium barbarum* | High | -2.94205037 |

| | | |
|---|---|---|
| *Leucobryum glaucum* | High | -2.924310682 |
| *Solanum sisymbriifolium* | High | -2.907547946 |
| *Cistus inflatus* | High | -2.906919411 |
| *Canna sp.* | High | -2.894583605 |
| *Ipomoea pubescens* | High | -2.889996201 |
| *Gleditsia sinensis* | High | -2.879039689 |
| *Solanum dulcamara* | High | -2.872083035 |
| *Kadsura heteroclita* | High | -2.871730779 |
| *Staurodesmus omearii* | High | -2.864594192 |
| *Salix eriocephala* | High | -2.858480138 |
| *Platanus occidentalis* | High | -2.856445994 |
| *Solanum ptychanthum* | High | -2.816659708 |
| *Schlegelia violacea* | High | -2.811023308 |
| *Passiflora edulis* | High | -2.806563029 |
| *Apios americana* | High | -2.798601132 |
| *Brodiaea sierrae* | High | -2.79100931 |
| *Brugmansia sanguinea* | High | -2.781428621 |
| *Xanthocercis zambesiaca* | High | -2.741392415 |
| *Calycanthus floridus* | High | -2.735882695 |
| *Salix sachalinensis* | High | -2.729762278 |
| *Aulacomnium heterostichum* | High | -2.72265933 |

| | | |
|---|---|---|
| *Astragalus propinquus* | High | -2.718076357 |
| *Datura metel* | High | -2.709371629 |
| *Gymnocladus dioicus* | High | -2.70773137 |
| *Buddleja lindleyana* | High | -2.707377545 |
| *Vittaria lineata* | High | -2.701351865 |
| *Rhodiola rosea* | High | -2.696009908 |
| *Racomitrium varium* | High | -2.686770936 |
| *Rehmannia glutinosa* | High | -2.682246431 |
| *Buddleja sp.* | High | -2.670018631 |
| *Hakea drupacea* | High | -2.649805752 |
| *Gleditsia triacanthos* | High | -2.647994052 |
| *Adiantum raddianum* | High | -2.633719335 |
| *Polyscias fruticosa* | High | -2.630388192 |
| *Akebia trifoliata* | High | -2.628419604 |
| *Nelumbo sp.* | High | -2.620258934 |
| *Bituminaria bituminosa* | High | -2.618059288 |
| *Impatiens balsamifera* | High | -2.616239267 |
| *Physcomitrium sp.* | High | -2.613837068 |
| *Copaifera officinalis* | High | -2.608001517 |
| *Hymenophyllum bivalve* | High | -2.601343128 |
| *Linum leonii* | High | -2.594147238 |
| *Physena madagascariensis* | High | -2.589981008 |
| *Ardisia humilis* | High | -2.577831861 |

| | | |
|---|---|---|
| *Schizolaena sp.* | High | -2.574176793 |
| *Cassytha filiformis* | High | -2.566569606 |
| *Santalum acuminatum* | High | -2.538580546 |
| *Ternstroemia gymnanthera* | High | -2.537724067 |
| *Grevillea robusta* | High | -2.524879038 |
| *Asparagus densiflorus* | High | -2.52321963 |
| *Jacquinia sp.* | High | -2.519971861 |
| *Aloe vera* | High | -2.519451993 |
| *Stemona tuberosa* | High | -2.517772275 |
| *Alangium chinense* | High | -2.499426548 |
| *Amaranthus tricolor* | High | -2.482969079 |
| *Ilex vomitoria* | High | -2.474138355 |
| *Hydrangea quercifolia* | High | -2.474011561 |
| *Pittosporum resiniferum* | High | -2.466777885 |
| *Gompholobium polymorphum* | High | -2.463700808 |
| *Gloriosa superba* | High | -2.462967888 |
| *Dichroa febrifuga* | High | -2.461623547 |
| *Illicium floridanum* | High | -2.447071519 |
| *Marattia attenuata* | High | -2.445113378 |
| *Ilex sp.* | High | -2.440160507 |
| *Tetrazygia bicolor* | High | -2.432977808 |
| *Bischofia javanica* | High | -2.428035903 |

| | | |
|---|---|---|
| *Linum lewisii* | High | -2.42287685 |
| *Plagiomnium insigne* | High | -2.422227995 |
| *Lycopodiella appressa* | High | -2.421574341 |
| *Roridula gorgonias* | High | -2.413867702 |
| *Senna hebecarpa* | High | -2.411958152 |
| *Daenikera sp.* | High | -2.409131357 |
| *Francoa appendiculata* | High | -2.408714522 |
| *Euptelea pleiosperma* | High | -2.407265828 |
| *Synsepalum dulcificum* | High | -2.394688978 |
| *Nepenthes alata* | High | -2.393261605 |
| *Meliosma cuneifolia* | High | -2.387924749 |
| *Rhododendron scopulorum* | High | -2.384212212 |
| *Amaranthus retroflexus* | High | -2.377962756 |
| *Sideroxylon reclinatum* | High | -2.360409833 |
| *Manilkara zapota* | High | -2.354440918 |
| *Myodocarpus sp.* | High | -2.352810014 |
| *Dicranum scoparium* | High | -2.352768818 |
| *Pittosporum sahnianum* | High | -2.3497829 |
| *Gyrostemon ramulosus* | High | -2.343265042 |
| *Glycyrrhiza lepidota* | High | -2.329062224 |
| *Nyssa ogeche* | High | -2.307044876 |

| | | |
|---|---|---|
| *Phytolacca americana* | High | -2.26168079 |
| *Symplocos tinctoria* | High | -2.25280139 |
| *Philonotis fontana* | High | -2.235219574 |
| *Timmia austriaca* | High | -2.211054487 |
| *Sassafras albidum* | High | -2.191605452 |
| *Persea borbonia* | High | -2.187350255 |
| *Glycyrrhiza glabra* | High | -2.163093611 |
| *Phytolacca bogotensis* | High | -2.124030485 |
| *Lagerstroemia indica* | High | -2.119003636 |
| *Cinnamomum camphora* | High | -2.109920077 |
| *Calocedrus decurrens* | High | -2.092855184 |
| *Acorus americanus* | High | -2.078546661 |
| *Encalypta streptocarpa* | High | -2.070541894 |
| *Cornus florida* | High | -2.038179507 |
| *Tetraclinis sp.* | High | -2.031219645 |
| *Ardisia revoluta* | High | -2.015768606 |
| *Cochlearia officinalis* | High | -1.999882646 |
| *Phormium tenax* | High | -1.990298274 |
| *Ochna serrulata* | High | -1.989198686 |
| *Sphagnum lescurii* | High | -1.974525571 |
| *Helonias bullata* | High | -1.968858741 |

| | | |
|---|---|---|
| *Ochna mossambicensis* | High | -1.965685789 |
| *Aucuba japonica* | High | -1.963297829 |
| *Sphagnum recurvum* | High | -1.932801357 |
| *Linum strictum* | High | -1.920457917 |
| *Freycinetia multiflora* | High | -1.912289703 |
| *Huperzia myrsinites* | High | -1.897225768 |
| *Reseda odorata* | High | -1.893738994 |
| *Sphagnum palustre* | High | -1.882035393 |
| *Huperzia squarrosa* | High | -1.855130071 |
| *Lonchitis hirsuta* | High | -1.845796959 |
| *Garcinia oblongifolia* | High | -1.839013144 |
| *Flaveria vaginata* | High | -1.832931758 |
| *Flaveria palmeri* | High | -1.827121547 |
| *Scouleria aquatica* | High | -1.821843148 |
| *Angelica archangelica* | High | -1.74893108 |
| *Papuacedrus papuana* | High | -1.739832784 |
| *Xerophyllum asphodeloides* | High | -1.738425415 |
| *Xanthium strumarium* | High | -1.732961732 |
| *Allionia incarnata* | High | -1.726161209 |
| *Allionia incarnata* | High | -1.722931868 |
| *Oxera neriifolia* | High | -1.697527136 |

| | | |
|---|---|---|
| *Fokienia hodginsii* | High | -1.625911848 |
| *Flaveria sonorensis* | High | -1.623336062 |
| *Pinus jeffreyi* | High | -1.579223884 |
| *Flaveria kochiana* | High | -1.578676215 |
| *Chamaecyparis lawsoniana* | High | -1.57627915 |
| *Pseudotsuga wilsoniana* | High | -1.483743248 |
| *Smilax bona-nox* | High | -1.482810719 |
| *Laurelia sempervirens* | High | -1.463538541 |
| *Microbiota decussata* | High | -1.401561799 |
| *Pinus radiata* | High | -1.376446242 |
| *Metasequoia glyptostroboides* | High | -1.366286201 |
| *Pinus ponderosa* | High | -1.364524049 |
| *Pinus parviflora* | High | -1.340843655 |
| *Planophila laetevirens* | High | -1.035502127 |

| Table S2. Full list of SpecKS configurable parameters | | | |
|---|---|---|---|
| **Category** | **Parameter/Tag** | **Default** | **Description** |
| Paths | output_folder_root | /home/SpecKS_output | Output destination. Write access required. SpecKS will postpend a date and timestamp to the directory name, as in “SpecKS_output_m04d26y2024_h12m21s55” so that subsequent runs are not overwritten. |
| Polyploid | name | [Auto/Allo]polyploid_N" | Name of polyploid. We suggest something descriptive. |
| Polyploid | DIV_time_MYA | 0 MY | DIV time in MY. Time of subgenome divergence. For gradual parental speciation, this will be the mode of the gene tree divergence times. |
| Polyploid | WGD_time_MYA | 0 MY | WGD time in MY. This will be the start time of ohnolog shedding, which will continue until the present time. |
| Polyploid | gene_div_time_distribution_parameters | impulse,1,1 | Distribution of divergence times for gene trees at DIV time. Currently-available choices are the Dirac Delta, exponential and lognormal distributions. For Dirac Delta use “impulse,1,1”. For exponentia, use the format “expon,0,K”, when K is the exponential decay constant. (We suggest Ne*Gt.). Lognormal distributions are also supported, with the format "lognorm,shape_parameter,xscale" (see scipy.stats, “lognorm” and “expon” for more details). For polyploids whose gene tree divergence might be a mix of distributions (ie, segmental allopolyploids), multiple distributions may be given, with the last parameter being the proportion of genes which belong in each distribution. Ie,<br><gene_div_time_distribution_parameters><br>impulse,1,1,0.5<br></gene_div_time_distribution_parameters><br><gene_div_time_distribution_parameters><br>expon,0,10,0.5<br></gene_div_time_distribution_parameters> |
| Species | full_sim_time | 100 MY | The length of the time period to simulate. |

| | | | |
|---|---|---|---|
| Tree | | | Note that since speciation is a gradual process, it may be necessary to start the simulation well in advance of the SPC time. |
| Gene Tree | mean_gene_birth_rate_GpMY | 0.001359 genes per MY | Gene birth rate. Note this may be lineage specific. Our default is chosen from [Guo 2013] |
| Gene Tree | SSD_half_life_MY | 4 MY | Half life of small-scale duplications. Our default is chosen from [Lynch and Conery 2003] |
| Gene Tree | WGD_half_life_MY | 31 MY | Half life of ohnologs (whole-genome duplications). Our default is based on [Guo 2013 and Maere 2005] |
| Gene Tree | num_gene_trees_per_species_tree | 3000 | Default is set to give a well-supported histogram without taking too long to run. If set too low, the final histogram will look too sparse. Higher numbers may be more realistic [Sterk 2007] and result in smoother histograms, but have a longer run time. Since gene trees are simulated independently, the number does not affect the general histogram shape, merely the number of samples in it, so high numbers are not always necessary. |
| Sequence Evolution | num_replicates_per_gene_tree | 1 | SpecKS can automatically run replicates for a given simulation, randomizing appropriately. |
| Sequence Evolution | num_codons | 1000 | The number of codons in each gene to be simulated. All genes in the sim have the same length. 1000 was chosen in agreement with [Tiley 2018]. |
| Sequence Evolution | Ks_per_Myr | 0.01 | K*s* per million years. This number is lineage and gene family specific, and may need to change depending on user needs [Gaut 1996 and Koch 2000] . The default of 0.01 was chosen in agreement with [Tiley 2018], and in range with[ Blanc and Wolfe 2004] |
| Sequence Evolution | per_site_evolutionary_distance | 0.01268182 | The per-site-evoultionary_distance is used to calculate the total tree length per gene tree input to evolver. The default setting was derived by [Tiley 2018] for a Ks_per_Myr of 0.01 and the evolutionary GY94 model. |

| | | | |
|---|---|---|---|
| Sequence Evolution | evolver_random_seed | 137 | The random seed used by evolver. We have exposed this so that the user can set this (or randomize it) directly as desired. The default value was simply chosen at random. |
| Main | LogFileName | log.txt | The name of the log file. The option to override the log file name supports the use case where the user might have multiple simultaneously executing runs sharing an output folder, and wishes them each to have a unique log file, to prevent overwriting or deadlock. |
| Histogram | maxKs | 5 | Debugging parameter. These histograms are only meant to give a confirmation of the run success. |
| Main | StopAtStep | 999 | Debugging parameter. If you want the sim to stop after only running the N'th module, set to N. |
| Main | IncludeVisuals | FALSE | Debugging parameter. If set to TRUE, .png files are generated for each gene tree at each stage. This can take up a lot of space on disk. |

**Table S3. Example SpecKS xml input file**

```xml
<metadata>
    <Paths>
        <output_folder_root>/usr/scratch2/userdata</output_folder_root>
    </Paths>
    <SpeciesTree>
        <polyploid>
            <name>MyAlloPolyploid</name>
            <DIV_time_MYA>20</DIV_time_MYA>
            <WGD_time_MYA>18</WGD_time_MYA>
            <gene_div_time_distribution_parameters>expon,0,1.6
            </gene_div_time_distribution_parameters>
        </polyploid>
        <polyploid>
            <name>MySegmentalPolyploid</name>
            <DIV_time_MYA>40</DIV_time_MYA>
            <WGD_time_MYA>38</WGD_time_MYA>
            <gene_div_time_distribution_parameters>expon,0,5,0.3
            </gene_div_time_distribution_parameters>
            <gene_div_time_distribution_parameters>impulse,1,1,0.7
            </gene_div_time_distribution_parameters>
        </polyploid>
        <full_sim_time>50</full_sim_time>
    </SpeciesTree>
    <GeneTree>
        <mean_gene_birth_rate_GpMY>0.07</mean_gene_birth_rate_GpMY>
        <SSD_half_life_MY>4</SSD_half_life_MY>
        <WGD_half_life_MY>31</WGD_half_life_MY>
        <num_gene_trees_per_species_tree>26000</num_gene_trees_per_species_tree>
    </GeneTree>
    <SequenceEvolution>
        <num_replicates_per_gene_tree>3</num_replicates_per_gene_tree>
        <num_codons>1000</num_codons>
        <Ks_per_Myr>0.01</Ks_per_Myr>
        <per_site_evolutionary_distance>0.01268182
        </per_site_evolutionary_distance>
    </SequenceEvolution>
    <Histogram>
        <maxKs>2</maxKs>
    </Histogram>
    <StopAtStep>999</StopAtStep>
    <LogFileName>SpecKS_log.txt</LogFileName>
    <IncludeVisuals>FALSE</IncludeVisuals>
</metadata>
```

| Table S4. Parameters used to match experimental observations | | |
|---|---|---|
| **Species** | **Parameter** | **Setting** |
| *Coffea arabica (sim 20)* | DIV_time_MY | 0.36 MY |
| | WGD_time_MYA | 0.30 MY |
| | gene_div_time_distribution_parameters | expon,0,1 |
| | mean_gene_birth_rate_GpMY | 0.175 |
| | SSD_half_life_MY | 5.2 MY |
| | WGD_half_life_MY | 4 MY |
| | Ks_per_Myr | 0.0250 |
| | per_site_evolutionary_distance | 0.03170455 |
| *Populus trichocarpa (sim 23)* | DIV_time_MY | 56 MY |
| | WGD_time_MYA | 56 MY |
| | gene_div_time_distribution_parameters | expon,0,28.6 |
| | mean_gene_birth_rate_GpMY | 0.01 |
| | SSD_half_life_MY | 5.7 MY |
| | WGD_half_life_MY | 17.14 MY |
| | Ks_per_Myr | 0.0035 |
| | per_site_evolutionary_distance | 0.004438637 |
| *Zea mays (sim 21)* | DIV_time_MY | 20 MY |
| | WGD_time_MYA | 14 MY |

| | | |
|---|---|---|
| | gene_div_time_distribution_parameters | expon,0,9 |
| | mean_gene_birth_rate_GpMY | 0.160 |
| | SSD_half_life_MY | 6.153 MY |
| | WGD_half_life_MY | 7.7 MY |
| | Ks_per_Myr | 0.0065 |
| | per_site_evolutionary_distance | 0.008243183 |
| Shared parameters | Num_gene_trees_per_species_tree | 20000 |
| | Full_sim_time | 30 MY |

| Table S5. SpecKS inferred parameters based on transcriptomic data | | | | | |
|---|---|---|---|---|---|
| | SpecKS Input | | SpecKS inferred | Reported | |
| Species | Ks per MY | DIV/ WGD time (in MY) | DIV/WGD time (in Ks) | DIV/WGD (MYA) | Reference |
| *Coffea arabica* | 0.0150 | 0.36/0.30 | 0.009/0.005 | ~0.046-0.66 | Salojarvi et al. 2023 |
| *Populus trichocarpa* | 0.0033 | 56/56 | 0.195/0.195 | ~58/58 | Dai et al. 2014 |
| *Zea mays* | 0.0065 | 20/14 | 0.13/0.12 | ~20.5/11.4 | Gaut and Doebley 1997 |
